# Metabolic engineering of *Escherichia coli* to modulate hydrogen sulfide levels in the mammalian gut

**DOI:** 10.64898/2026.07.23.738723

**Authors:** Justin A. Hayes, Brent Buchinger, Anna W. Lunger, Tanvi Khanvilkar, William Gasparrini, Matthew T. Fernez, Madeleine Morrissette, Philip Strandwitz, Abigail N. Koppes, Ryan Koppes, Benjamin M. Woolston

**Affiliations:** Department of Chemical Engineering, Northeastern University, 360 Huntington Ave. Boston, MA 02115, USA; Holobiome Inc., 650 Albany St., Boston, MA 02118, USA; Department of Bioengineering, Northeastern University, 360 Huntington Ave. Boston, MA 02115, USA; Department of Chemistry and Chemical Biology, Northeastern University, 360 Huntington Ave., Boston, MA 02115, USA

**Keywords:** metabolic engineering, host-microbe interactions, hydrogen sulfide

## Abstract

Hydrogen sulfide (H_2_S) is a microbiota-derived metabolite in the gastrointestinal tract implicated in a number of diseases. Its volatility and reactivity make experimentally controlling H_2_S concentration *in vivo* difficult, limiting our ability to interrogate its dose-dependent effects on host physiology. Engineered bacteria present a compelling solution, yet most probiotic metabolic engineering approaches have focused on *in vitro* optimization, failing to account for the complex intestinal environment. Here, we engineered *Escherichia coli* strains to produce or consume H_2_S in specific intestinal regions by incorporating knowledge of the local metabolic environment and resident microbial activities into the design process. Analysis of human-derived *ex vivo* cultures revealed that glutathione (GSH) is inefficiently converted to H_2_S, suggesting GSH as a relatively stable substrate for engineered sulfide production. We thus engineered a GSH-dependent H_2_S producer, which increased levels 21-fold *ex vivo*. To target the nutrient-rich, microbially sparse environment of the small intestine, we optimized a H_2_S producer that uses L-cysteine as a sulfur source, demonstrating a 7-fold increase in H_2_S levels in mice. Finally, to develop strains capable of sequestering H_2_S, we leveraged the availability of fumarate and nitrate as electron acceptors in the large intestine by engineering a strain expressing sulfide:quinone oxidoreductase (Sqr). This enables oxidation of H_2_S to intracellular polysulfides and achieves higher consumption rates than alternative sequestration strategies reliant on resource-intensive GSH production. Together, this work developed engineered microbes as precision tools to modulate H_2_S levels and showcases a generalizable framework for region-targeted design of engineered probiotics.

## 1. Introduction

Gut microbes influence host physiology in numerous ways, through degradation of dietary components, production of metabolites, and stimulation of host receptors^1–3^. Hydrogen sulfide (H_2_S) is a microbiota-derived metabolite whose concentration-dependent effects on host physiology include regulation of inflammation, vasodilation, and neurotransmission^4–7^. It has also long been associated with gastrointestinal disorders such as inflammatory bowel disease (IBD) and colorectal cancer, although its mechanistic role in disease is widely debated, with evidence supporting both a pro- and anti-inflammatory role. For example, H_2_S has been shown to promote ulcer healing and exert anti-inflammatory effects in rodents and humans^8–10^. In contrast, the addition of H_2_S-producing microbes can induce an IBD-like phenotype in animals^11^. Building on this preclinical study, ulcerative colitis patients present with increased fecal H_2_S levels that correlate with disease severity, and patients with irritable bowel syndrome diarrhea subtype exhibit significantly higher breath sulfide^12–14^. As a volatile gasotransmitter with a pKa_1_ near physiological pH, sulfide exists largely in the gaseous form in vivo, making it difficult to handle, titrate, and precisely control in experimental or clinical settings^15–17^. Conflicting reports on the role of microbially derived H_2_S partially reflect the inability to control local sulfide concentration and exposure duration.

Most studies evaluating the impact of H_2_S have used chemical donors, but these offer limited control over sulfide concentrations in the gastrointestinal (GI) tract. Some release sulfide rapidly (e.g., Na_2_S, NaHS), exhibit pH-dependent release (GYY4137), or require reducing agents (allyl polysulfides) for activity^18^. Biomaterial-based delivery systems also struggle to achieve precise dosing due to pH, redox, and water content differences *in vivo*^16^. These challenges are exacerbated by regional heterogeneity of the chemical and metabolic environment along the intestinal tract^15^. Precisely modulating intestinal H_2_S remains a major challenge that impedes efforts to understand its mechanistic role in disease^19,20^.

Engineered bacteria offer a powerful platform for modulating intestinal metabolite levels for diagnostic, therapeutic and research purposes^7,21–23^. Several engineered probiotics have been tested clinically, including a strain engineered for intestinal phenylalanine degradation for phenylketonuria patients^20^. Another example is a strain engineered to metabolize a unique dietary polysaccharide unavailable to other gut microbes, allowing robust colonization at high relative abundance. The colonizing strain was further engineered to degrade oxalate in patients with hyperoxaluria^23^. Another group identified arginine:glycine amidinotransferases from native gut microbes and engineered a strain to synthesize guanidinoacetic acid from arginine and glycine, which is then converted by the liver to creatine to improve cognitive performance^23^. We previously engineered bacteria to titrate H_2_S levels in human gut-on-chip systems, achieving lower variability than a standard sulfide donor^7^. Together, these studies highlight the potential of engineered microbes to modulate gut metabolites, motivating targeted strategies to control intestinal sulfur metabolism *in vivo*.

There are several well characterized pathways that could be leveraged in engineering a probiotic to produce or sequester sulfide. Many gut microbes respire inorganic sulfur, such as sulfate, sulfite, and thiosulfate, through dissimilatory reduction with H_2_S as a by-product^25^. Similarly, sulfur-containing amino acids such as L-cysteine and taurine can be fermented to H_2_S. In a metatranscriptomic analysis of 318 individuals, cysteine-degrading genes, including cysteine desulfhydrases, desulfurases, desulfidases, and aminotransferases, were expressed in 100% of samples^26^. Conversely, the microbiota can also reduce H_2_S levels through L-cysteine biosynthesis or oxidation via sulfide quinone reductase (*Sqr*)^27,28^. Together, these pathways provide multiple possible options for metabolic engineering to enable localized, tunable, and sustained enzymatic control of H_2_S levels *in vivo*.

Beyond identifying the pathway for producing or sequestering the metabolite of interest, probiotic design must consider core metabolic engineering principles, which can affect engineered strain functionality *in vivo*. For example, Novome Biotechnologies developed an engineered strain of *Bacteroides vulgatus* expressing a five-gene oxalate-degradation pathway as a therapeutic for kidney stones. Although the pathway was functional, the engineered strain exhibited reduced growth rate *in vitro* in the presence of oxalate, indicating that oxalate degradation imposed a fitness cost. Although the strain successfully colonized healthy humans with low oxalate levels, the pathway was lost in patients with oxalate-associated disease^21^. Colonization with a strain carrying a burdensome pathway led to pathway loss at the population level in patients requiring treatment, highlighting the need to consider selective pressures *in vivo* in the overall engineering strategy.

Additionally, the heterogeneous metabolic environment along the GI tract creates region-specific nutrient profiles that could influence engineered pathway function. The small intestine is characterized by abundant dietary peptides and nutrients, mildly acidic pH (5-6), relatively high oxygen tension, and sparse microbial populations, with nearly 90% of dietary nutrient absorption occurring in this region^29^. In contrast, the colon exhibits limited dietary substrate availability, neutral to slightly alkaline pH, anaerobic conditions, and dense microbial communities exceeding 10^11^ cells per gram of luminal content^29^. Native gut microbes colonize discrete intestinal regions defined by local metabolic niches, defined by differentially available carbon sources, oxygen tension, microbial competition, and host factors such as mucus. As a result, most taxa do not occupy the full length of the gastrointestinal tract, and their metabolic activity is inherently spatially constrained. These regional differences impose distinct selective pressures and metabolic constraints that must be accounted for when engineering bacteria for targeted intestinal delivery.

Here, we developed a panel of engineered *Escherichia coli* (*E. coli*) strains to modulate intestinal H_2_S levels. In the process, we studied fecal communities from five human donors and uncovered insights into sulfur metabolism of the gut microbiota. We integrated these findings with principles of gastrointestinal physiology to engineer strains to target either the small or large intestine. The strains were designed to function within each intestinal region, accounting for variations in nutrient availability, competition for sulfur sources, oxygen tension, and metabolic competition from resident microbiota. These engineered bacteria serve dual purposes: 1) as research tools to interrogate host-microbiome H_2_S interactions and 2) as medical solutions for gastrointestinal diseases characterized by dysregulated H_2_S levels. More broadly, this work establishes generalizable design principles and a workflow for developing robust engineered probiotics tailored to the complexities of the intestinal metabolic environment.

## 2. Methods

### 2.1 Quantification of H_2_S and thiols

#### 2.1.1 Methylene blue assay for H_2_S quantification

200 µL of the experimental sample was added to 615 µL zinc acetate mixture (600 µL of 1% w/v zinc acetate dihydrate with 15 µL 3M NaOH) and vortexed. After 5-10 min of incubating at room temperature, 150 µL of 0.1% N,N-dimethyl-p-phenylenediamine (DMPD) in 5M HCl was added, followed by 150 µL of 23 mM FeCl_3_ in 1M HCl. Samples were centrifuged at 16,000 RCF for 2 min. 200 µL of supernatant were plated in a transparent 96-well plate and were quantified using absorbance (670 nm) and appropriate standard curve made with Na_2_S.

#### 2.1.2 Quantification of sulfur metabolites with monobromobimane and HPLC (*in vitro* experiments)

100 µL of experimental sample was centrifuged at 16,000 RCF and the supernatant was stored at -80C until use. 7 µL of sample was mixed with 2 µL 1-octanol and 3 µL of 6M sodium borohydride:DMSO (2:1 volume ratio) mixture for 15 minutes at room temperature to reduce oxidized thiols (L-cysteine and glutathione) in the sample. 0.6 µL of 3M HCl was added to terminate the reduction process. 90 µL 3 mM monobromobimane (mBBr) in 200 mM Tris-HCl (pH 9.5) and 200 µM ethylenediaminetetraacetic acid (EDTA, prepared as 200 mM in 3 M NaOH) was mixed with the sample, covered in foil, and incubated at room temperature for two hours to derivatize the thiols. After incubation, 30 µL 1M HCl was added to terminate the derivatization reaction, samples were centrifuged to remove debris, and 100 µL supernatant was added to HPLC vials.

#### 2.1.3 Hot cyanolysis for polysulfide quantification

Sqr activity was measured using a modified cyanolysis-based assay adapted from prior work^30^. Briefly, 250 μL of each sample was combined with 550 μL of 1% (w/v) boric acid and 200 μL of 100 mM potassium cyanide in microcentrifuge tubes. Mixtures were heated to 100C for 5 minutes, then allowed to cool to ambient temperature. After cooling, 100 μL of ferric nitrate solution was prepared by dissolving 3 grams of ferric nitrate in 5 mL of 33% perchloric acid. Samples were centrifuged at 16,000 g for 5 minutes, and 200 μL of the supernatant was transferred to a 96-well plate. Absorbance was measured at 460 nm. Sulfur species concentrations were determined by comparison to a sodium thiosulfate standard curve processed in parallel, with standards supplemented with 5 μL of 1M copper sulfate to catalyze thiosulfate conversion.

#### 2.1.4 Quantification of intestinal sulfur metabolites with monobromobimane and HPLC-FLD

The mBBr assay was adapted to detect thiol compounds in the gut luminal content. 1.5 mL centrifuge tubes containing 250 μL of 3 mM mBBr dissolved in 200 mM Tris-HCl (pH 9.5) with 200 μM EDTA were weighed before sampling. Intestinal luminal content was sampled from three locations (most proximal, mid-point, and most distal) for both the stomach and small intestine, and one location of the cecum and colon. Approximately 10-20mg of content was placed into a 1.5 mL centrifuge tube containing mBBr solution. Samples were immediately weighed and then centrifuged at 21,000 RCF for 1 minute. 200 μL of supernatant was added to clean 1.5 mL centrifuge tubes for 1.5 hours at room temperature covered in aluminum foil. 60 μL of 1M HCl was added to terminate the reaction. Samples were centrifuged to remove excess debris, and supernatant filtered through 0.2 µm filters into HPLC vials. Quantification was done using the HPLC method and standard curve as described above. Values were normalized to sample weight (concentration per unit mass).

An Agilent 1260 HPLC Infinity II system was used with a gradient elution method consisting of mobile phase A (10% methanol, 89.75% water, 0.25% acetic acid, pH 3.9) and mobile phase B (89.75% methanol, 10% water, 0.25% acetic acid, pH 3.9) at 0.5 mL/min. The gradient applied to separate thiols: 0-2 min, 0% B; 2-8 min, 46% B; 8-9 min, 64% B; 9-13 min, 100% B; 13-14 min, 0% B; 14-15 min, 0% B. 10 uL of sample was injected into InfinityLab Poroshell 120 EC-C18, 3.0 x 100 mm, 2.7 µm C18 reversed phase column at 40C. Compounds were identified with a fluorescence detector. Calibration curves of the pure compounds were used to identify and quantify compounds in experimental samples.

### 2.2 Strain Engineering

#### 2.2.1 Cell engineering, gene knockouts, and plasmid assembly

All microbial strains used are derivatives of *Escherichia coli* K-12 MG1655 (DE3) or S1030. Genes were deleted from the genome using lambda-red recombination and CRISPR Cas9 systems following previously published protocols^31^. Plasmids were assembled using polymerase chain reaction (PCR) and Gibson Assembly techniques. All plasmids were chemically transformed into the respective *E. coli* strain and used for experiments. All plasmids, primers, and strains are listed in **Table S1, S2, and S3.**

Gene deletions of *cyuR*, *iscS*, *cyuA*, and *cyuP* were generated in *E. coli* MG1655 using a combination of λ-Red recombineering and CRISPR-Cas9-mediated genome editing^31^. The CRISPR components were derived from the plasmids pCas (Addgene #62225) and pTargetF (Addgene #62226). For each target gene, a gene-specific 20-nucleotide guide sequence was introduced into pTargetF by PCR. Cells were sequentially transformed with pCas and the corresponding pTargetF construct, followed by plasmid curing as described^31^. Colonies were screened by PCR, and correct genomic edits were confirmed by Sanger sequencing (Genewiz, Azenta Life Sciences).

*E. coli* genes (*ggt*, *pepT*, *pepB, cyuA, cyuP, gshA, gshB*) were cloned from gDNA extracted from *E. coli* MG1655(DE3) with a Monarch gDNA Spin Kit (#T3010, New England Biolabs Inc.) via PCR. The *fupA* gene (from *Francisella tularensis*) and *gshF* (from *Streptococcus thermophilus*) were codon harmonized for *E. coli* and synthesized (Integrated DNA Technologies) (**Table S4**). The *sqr* constructs were previously assembled in our lab^32^.

DNA constructs were generated using Gibson Assembly (New England Biolabs, E2621) to join plasmid backbones with genes of interest. Assembly reactions were introduced into chemically competent *E. coli* DH5α cells (New England Biolabs, C2987) by transformation, and transformants were selected on antibiotic-containing agar plates. Individual colonies were isolated and screened by PCR, with correct constructs confirmed by Sanger sequencing. Verified plasmids were purified using a miniprep kit (Zymo Research, D4211) and subsequently transformed into the appropriate chemically competent production strains. A complete list of strain designations and descriptions is provided in **Table S1**. Primer sequences used for cloning are listed in **Table S2**, full plasmid maps are provided in **Table S3**, and DNA sequences in **Table S4**.

#### 2.2.2 Growth curve experiments

Cultures were initiated by overnight growth in LB medium. Cells were harvested by centrifugation, the spent medium was discarded, and pellets were resuspended in either LB or M9 minimal medium supplemented with glucose (4 g/L), casamino acids (1 g/L; VWR, 2240), and thiamine (1 mg/L; Sigma-Aldrich, T4625). Resuspended cultures were diluted to an initial OD_600_ of 0.01 and dispensed into 96-well plates covered with a plastic lid (Fisher, 12565501). Growth was monitored for 16 hours at 37 °C using a plate reader (SpectraMax i3, Molecular Devices), with continuous shaking at medium speed and optical density measurements recorded at 10-minute intervals. All conditions were run in technical triplicate.

### 2.3 *in vitro* H_2_S experiments

#### 2.3.1 Cell growth and experimental preparation

L-cysteine H_2_S-producing strain: Precultures from cryostocks were inoculated in LB media and grown aerobically overnight in a 37°C shake incubator (200 RPM). 300 μL of overnight culture was inoculated in 15 mL LB media in an aerated 125 mL Erlenmeyer flask and put in a 4°C static incubator overnight. In the morning, the temperature was automatically adjusted to 37°C and shaking activated. After 2.5 hours of growth, cells were induced with 100 μM isopropyl β-d-1-thiogalactopyranoside (IPTG) and grown for an additional two hours.

GSH H_2_S-producing strain: Cryostocks were revived as above. 300 μL of overnight culture was inoculated in 15 mL LB media in an aerated 125 mL Erlenmeyer flask and put in a 20°C shaking incubator (200 RPM) for 16 hours overnight, induced with 100 μM IPTG and 10 mM L-arabinose. It has been shown that *ggt* mRNA in *E. coli* is more stable at 20°C than 37°C and subsequently higher *ggt* activity^33^.

gshAB/gshF strains: Cryostocks were revived as above. 300 μL of overnight culture was inoculated in 15 mL LB media in an aerated 125 mL Erlenmeyer flask and put in a 20°C shaking incubator (200 RPM) for 16 hours overnight, induced with 100 μM IPTG. To counteract slow growth rates caused by genetic knockouts and low-temperature induction, the incubator was adjusted to 37°C for three hours to stimulate cell growth.

Sqr strain: Cryostocks were revived as above. 300 uL of overnight culture was inoculated in 15 mL LB media in a 125 mL Erlenmeyer flask and put in a 37C shake incubator (200RPM) for 2 hours. Then, 10 mM L-arabinose was added for 2 hours to induce gene expression.

#### 2.3.2 Hungate tube experiments

For aerobic experiments, induced cells were centrifuged, resuspended in experimental media to OD_600_ 0.5, and sealed in sterile Hungate tubes. For anaerobic experiments, induced cells were centrifuged and moved into the anaerobic chamber for handling. Cells were resuspended in anaerobic experimental media and sealed in sterile anaerobic Hungate tubes. Tubes were moved to a shaking incubator at 37°C (200 RPM). Samples were taken with sterile needles while maintaining a closed and sterile system. For anaerobic samples, the sterile needles and syringes were first flushed with pure nitrogen and used for sampling. OD_600_ was measured at time of inoculation and at the time of sampling, and metabolite measurements were normalized to OD_600_.

#### 2.3.3 Small intestinal content *ex vivo* experiments

Rats were sacrificed and small intestinal luminal content was mixed with sterile PBS (1g content per 5mL PBS) and homogenized, centrifuged, and supernatant used as a medium. 5mL of the supernatant were transferred to Hungate tubes and 1 mM L-cysteine, 1 mM GSH, or 2 mM Na_2_S were added, along with the appropriate antibiotic. Engineered bacteria or control bacteria were added to OD_600_ 0.5. After three and 20 hours, samples were taken and measured with the methylene blue, mBBr, or cyanolysis assays to quantify metabolites of interest.

#### 2.3.4 Human fecal consortia H_2_S metabolism experiments

Five human fecal samples were acquired from Holobiome (Advarra Institutional Review Board Protocol: Pro00069198). In an anaerobic chamber (Coy Labs), cryostocks were inoculated into anoxic brain heart infusion media in serum bottles with butyl rubber stoppers. Cultures were moved to a 37°C shake incubator (200 RPM) for overnight growth. After growth, the cultures were transferred to the anaerobic chamber, centrifuged at 16,000 RCF and resuspended in 5 mL of anaerobic minimal glucose media, M9, to OD_600_ 0.5. 1 mM of each sulfur substrate (L-cysteine, glutathione, sulfite, thiosulfate) was added and the cultures were placed in Hungate tubes to maintain anaerobic conditions and preserve H_2_S gas tension. For experiments testing H_2_S degradation, 2 mM sodium sulfide, 20 mM sodium nitrate, and or 20 mM sodium fumarate were added to the cultures. The cultures were moved to a 37°C shake incubator (200 RPM) and H_2_S was measured using the methylene blue assay at 0, 3, and 20 hours after incubation.

#### 2.3.5 Testing engineered bacteria in human fecal consortia

The fecal consortia and engineered cells were prepared as described above. After growth, the cultures were transferred to an anaerobic chamber, centrifuged at 16,000 RCF and resuspended in an anaerobic minimal glucose media, M9, supplemented with products for each experiment. For H_2_S production experiments, the cell density of the GSH strain and human consortia were equalized (1:1 cell ratio) at OD_600_ 0.5 in anaerobic M9 minimal media. 1 mM glutathione was spiked into the anaerobic culture and placed in Hungate tubes in a 37C shake incubator. The methylene blue assay was used to quantify sulfide.

For the *gshAB/gshF-*expressing strains, 1.5 mM of H_2_S, 2 mM L-serine, 20 mM glycine, and 20 mM glutamic acid were added and the anaerobic cultures were placed in Hungate tubes. For the *sqr*-expressing strain, 1.5 mM of H_2_S, 20 mM sodium nitrate, and or 20 mM sodium fumarate were added, and the cultures were placed in Hungate tubes. The cultures were moved to a 37°C shaking incubator (200 RPM) and H_2_S was measured using the methylene blue assay at 0, 3, and 20 hours after incubation. 100 μL of each culture was taken, centrifuged at 16,000 RCF, and supernatant was collected and stored at -80°C until quantification with monobromobimane and HPLC.

### 2.4 Animal experiments

#### 2.4.1 IACUC approvals and animal information

The use of animals was approved by the Northeastern University Institutional Animal Care and Use Committee (IACUC) (Protocol# 23-1122R-A2). Male and female Sprague-Dawley rats (150-175 grams), and male and female C57BL/6 mice aged 8-12 weeks from Charles River Laboratories or Jackson Laboratories were group-housed in a temperature-controlled environment and allowed to acclimate at the Northeastern University animal facilities for 72 h. Animals were provided food and water *ad libitum* and exposed to a 12-h light/dark cycle. Animals were randomly assigned to groups.

#### 2.4.2 Evaluation of H_2_S-producing microbial strains in C57BL/6 mice

L-cysteine-H_2_S or empty vector cells to be administered were grown up as described above. For experiments where there were two separate gavages: cells were concentrated to 2x10^10^ CFUs and resuspended in 120 mM sodium bicarbonate and gavaged to animals (120 μL). A separate solution was made of 120 mM sodium bicarbonate and 150 mM L-cysteine and gavaged (60 μL). 60 minutes post-gavage, animals were sacrificed and luminal contents sampled as described below. For experiments where there was one gavage: cells were concentrated to 2x10^10^ CFUs and resuspended in 170 mM sodium bicarbonate and 100 mM L-cysteine, and gavaged (total 120 μL).

#### 2.4.3 Evaluation of Sqr strain in C57BL/6 mice

Sqr cells to be administered were grown as described above. Cells were concentrated to 2x10^10^ CFUs and resuspended in 120 mM sodium bicarbonate and gavaged to animals (120 μL gavage). A separate solution was made of 120 mM sodium bicarbonate, 50 mM GYY4137, and 150 mM sodium fumarate (60 μL gavage). 60 minutes post-gavage, animals were sacrificed and luminal contents sampled as described below.

#### 2.4.4 Evaluation of H_2_S-producing and Sqr strains in C57BL/6 mice

L-cysteine-H_2_S and Sqr cells to be administered were grown up as described above. Both cell types were concentrated to 2x10^10^ CFUs and resuspended in 120 mM sodium bicarbonate and gavaged to animals (120 μL gavage). A separate solution was made of 120 mM sodium bicarbonate, 150 mM L-cysteine, and 150 mM sodium fumarate (60 μL gavaged). 60 minutes post-gavage, animals were sacrificed and luminal contents sampled as described below.

### 2.5 Statistical Analysis

Data were assessed for normality using the Shapiro-Wilk test. Normally distributed data were analyzed using Welch’s t-test for two-group comparisons, one-way ANOVA, or two-way ANOVA with Tukey’s post-hoc test for multiple group comparisons (*p < 0.05). Non-normally distributed data were analyzed using Kruskal-Wallis test with Dunn’s post-hoc test for multiple group comparisons. For non-normal data, when negative controls were included (i.e., absence of H_2_S production or consumption), analyses comparing treatments were conducted excluding the negative control to specifically assess statistical differences among treatments and avoid skewing from the large block of structural zeros. Statistical details of each experiment can be found in the figure legends. Bar graphs represent the average value from biological replicates, and dots represent the result from each biological replicate. Error bars represent one standard deviation. GraphPad Prism was used to carry out the analysis. Graphs were formatted in Adobe Illustrator.

## 3. Results

### 3.1. Understanding human fecal microbiota H_2_S metabolism

Though multiple substrates and pathways exist for the production of sulfide, their relative activities within the gastrointestinal tract have not been rigorously evaluated. We hypothesized that screening sulfide production capacity from different substrates in human-derived microbial consortia could identify promising substrates not readily metabolized by native microbes. We acquired five human fecal microbiota samples, then tested each consortia individually for their ability to produce H_2_S from 1 mM of four sulfur species under anaerobic conditions: L-cysteine, glutathione, sulfite, and thiosulfate (**Fig. 1a**). As expected, L-cysteine was metabolized rapidly to sulfide within three hours by all cultures, and with significantly greater conversion than all other substrates (86% ± 30% conversion of sulfur added, *p*<0.001). Sulfite (9% ± 12%), thiosulfate (5% ± 8%), and GSH (1% ± 4%) were poorly converted to H_2_S within the same 3-hr time frame. After 20 hours, sulfide production from GSH was significantly lower than from L-cysteine (27% ± 25% vs 70% ± 9%, *p*<0.001), sulfite (57% ± 30%, *p*<0.05), and thiosulfate (60% ± 19%, *p*<0.05, **Fig. 1b**). H_2_S conversions from each sulfur substrate sorted by each human donor can be found in **Supp. Fig. 1**. These data show each community’s ability to degrade each substrate, highlighting ‘responder’ and ‘non-responder’ communities. Of the substrates tested, GSH was the most stable, with minimal conversion to H_2_S across these human fecal microbiota samples.

**Fig. 1.**
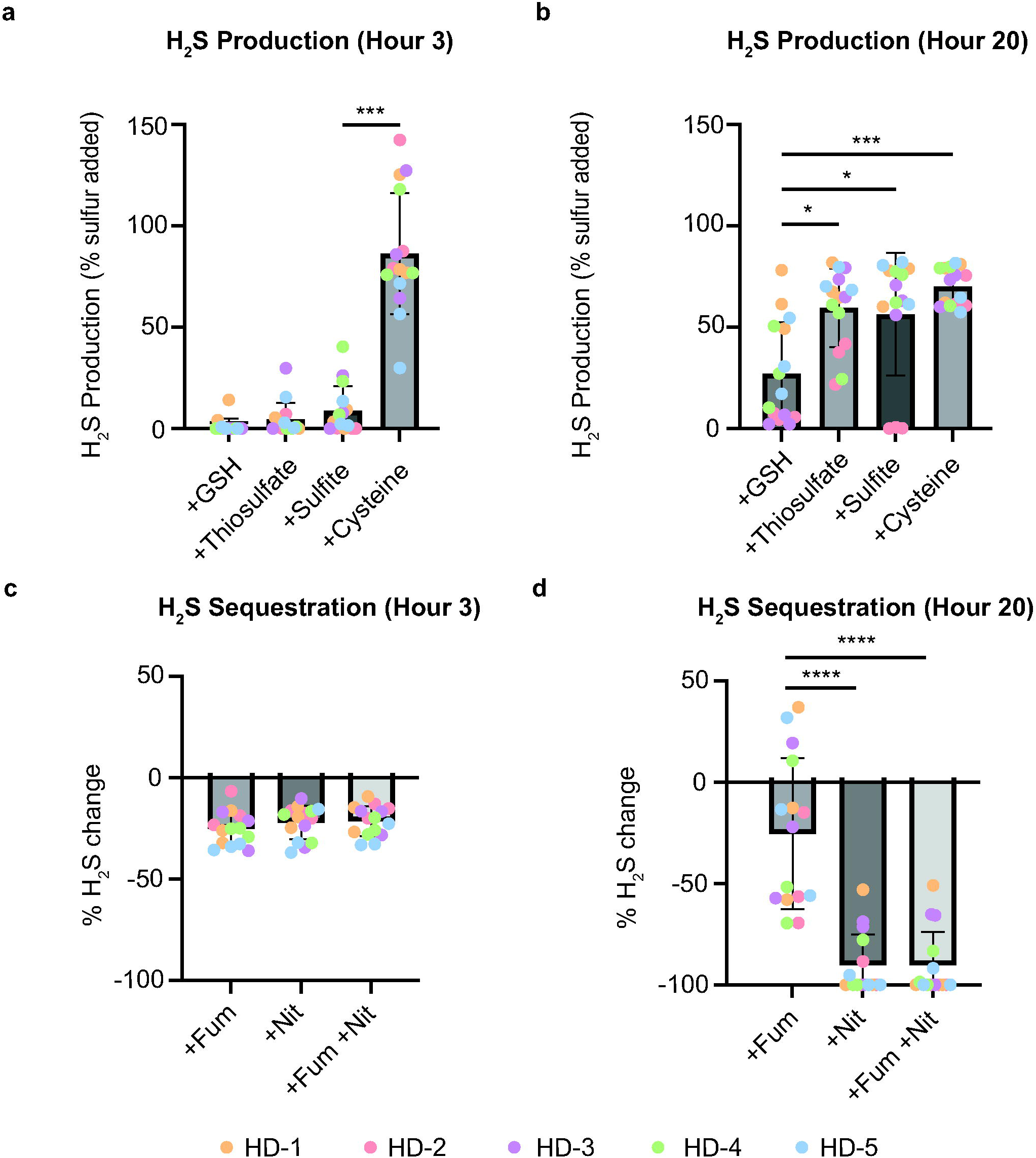
Human fecal cultures poorly convert glutathione to H_2_S and degrade H_2_S mediated by nitrate. Five human donor fecal cultures (HD1-5) were grown overnight anaerobically and resuspended in M9 minimal media and 1 mM of L-cysteine, glutathione, thiosulfate, or sulfite. Dot colors correspond to each human donor, see legend. Due to the presence of two sulfur atoms in thiosulfate, H_2_S production was normalized to the number of sulfur atoms in the substrate and reported as H_2_S per mol sulfur equivalents. H_2_S levels were measured via the methylene blue assay after **a)** 3 hours and **b)** 20 hours. Fecal consortia were prepared in the same manner and mixed with ∼1.5 mM Na_2_S with or without electron acceptors, fumarate (Fum) and nitrate (Nit). H_2_S levels were measured after **c)** 3 hours and **d)** 20 hours. n = 3 independent experiments. Error bars represent SD, and bars represent the mean value. *p<0.05, ∗∗p < 0.01, ∗∗∗p < 0.001, ∗∗∗∗p < 0.0001. Kruskal-Wallis test with Dunn’s multiple comparisons was used, with statistical significance set at p < 0.05.

We next evaluated the ability of the five fecal consortia to oxidize H_2_S, using the electron acceptors nitrate and fumarate under anaerobic conditions, two molecules commonly found in the gut lumen^34,35^. After three hours, there was no reduction in sulfide levels (**Fig. 1c**). After 20 hours, cultures supplemented with nitrate significantly reduced H_2_S levels relative to fumarate (-90% ± 15% vs. -25% ± 37%, *p*<0.0001, **Fig. 1d**). Based on these data, the five human samples couple sulfide oxidation to nitrate reduction, but not fumarate reduction. Sulfide quinone reductases (*Sqr*) oxidize H_2_S and have been found in the human gut microbiome, suggesting a possible mechanism for sulfide degradation^36^. These experiments approximate sulfide metabolism by the microbiota of the distal colon, an environment characterized by anoxic conditions, high microbial density, fecal-derived consortia, and the presence of alternative electron acceptors.

### 3.2 Engineered bacteria elevate upper intestinal H_2_S levels in mice

We previously developed a series of engineered bacteria to titrate H_2_S from L-cysteine in human gut-on-chip systems^7^. Here we set out to test these engineered H_2_S-producing strains *in vivo*. We benchmarked the engineered bacteria against the sulfide donor GYY4137, which has been used extensively as a slow-release sulfide donor for studying its role in biology^37,38^. We orally gavaged GYY4137 to mice and measured luminal H_2_S levels along the GI tract (**Fig. 2a**). After one hour, GYY4137 significantly increased stomach H_2_S compared to untreated animals (**Fig. 2b**). Small intestine, cecum, and colon levels remained unchanged (**Fig. 2c-e**).

**Fig. 2.**
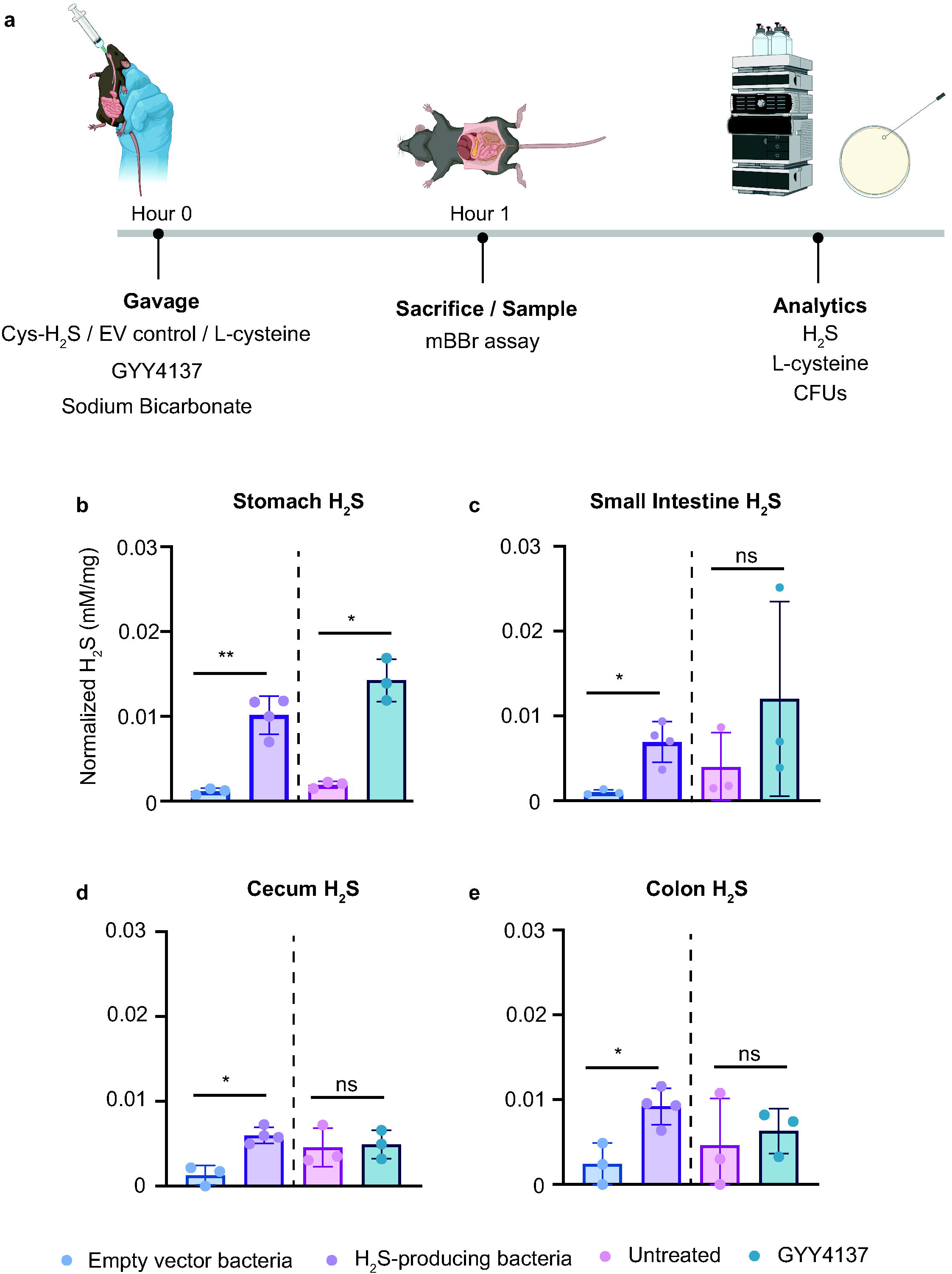
Engineered bacteria elevate upper intestinal H_2_S levels in mice. **a)** Animals were gavaged with GYY4137, L-cysteine-H_2_S-producing strain and L-cysteine, or empty vector control and L-cysteine. After 1 hour, animals were sacrificed and luminal content mixed with the mBBr assay. H_2_S values normalized to sample weight in the **b)** stomach **c)** small intestine, **d)** cecum, and **e)** colon. For no treatment control n=3, 2 females, 1 male; GYY4137: n=3, 2 females, 1 male. For EV control n=3, 2 female, 1 male; L-cysteine-H_2_S: n=4, 2 male, 2 female. Error bars represent SD, and bars represent the mean value. *p<0.05. Significance determined by unpaired t test with Welch’s correction, with statistical significance set at p < 0.05. Comparisons were only compared across respective controls. EV control and H2S-producing strains compared, GYY4137 compared to untreated animals. Figures made with Biorender.com

To deliver sulfide to the small intestine, where amino acids are abundant and competition from other microbes is limited, we used our previously developed strain that converts L-cysteine to H_2_S via overexpression of *cyuA* and *cyuP* (L-cysteine-H_2_S strain). To test its capacity to produce sulfide *in vivo*, we mixed small intestinal luminal content from rats with the L-cysteine-H_2_S strain or an empty vector control in the presence of L-cysteine. The engineered strain significantly elevated H_2_S levels within 3 hours, within the 2-8 hours window of small intestinal transit time (**Supp. Fig. 2**)^29^. Background sulfur and thiol content are identical across conditions, as each aliquot comes from the same homogenate and receives the same L-cysteine supplement. The only variable is expression of the H_2_S-producing genes. Any difference in H_2_S signal therefore reflects the activity of those enzymes. These data suggest this strain is well-suited to target the small intestine, leveraging the low abundance of other microbes (and thus limited competition for L-cysteine), sustained sulfide production in the presence of a luminal content matrix, and rapid reaction kinetics within the small intestinal transit window^29^.

The L-cysteine-H_2_S strain or an empty vector control were dosed to mice with L-cysteine as a sulfur substrate. After one hour, animals were sacrificed and intestinal content analyzed (**Fig. 2a**). The L-cysteine-H_2_S strain significantly elevated H_2_S 8.6-fold in the stomach (**Fig. 2b**) and 7-fold in the small intestine compared to control (**Fig. 2c**). Cecum levels were 4.6-fold higher (**Fig. 2d**) and 3.8-fold higher in the colon (**Fig. 2e**). Intestinal L-cysteine was also measured along the intestinal tract, peaking in the upper gut and rapidly approaching zero in the cecum and colon (**Supp. Fig. 3**). Notably, cecum and colonic L-cysteine levels were nearly undetectable, suggesting the L-cysteine is absorbed by the host and or converted to H_2_S in the upper GI tract by the engineered microbe.

### 3.3 Engineered bacteria for producing H_2_S from glutathione in the lower gut

Based on the relatively high stability of GSH in the fecal microbiota (**Fig. 1**), we engineered a strain to utilize GSH as a substrate for H_2_S production in the colon. Using GSH instead of other sulfur substrates could enable a high degree of control over colonic H_2_S production, as it is poorly converted to H_2_S by the native microbiota. We engineered a panel of strains overexpressing the *E. coli*-native glutathione hydrolase (*ggt*), peptidase T (*pepT*), peptidase B (*pepB*), L-cysteine transporter (*cyuP*), and L-cysteine desulfidase (*cyuA*). PepT and GGT degrade the tripeptide GSH into L-cysteinylglycine, followed by hydrolysis by PepB into L-cysteine. CyuP imports L-cysteine and CyuA degrades L-cysteine to H_2_S, which passively diffuses out of the cell (**Fig. 3a**). We previously showed overexpressing the native *E. coli* L-cysteine desulfidase *cyuA* and L-cysteine importer *cyuP* produced 53-fold more H_2_S from L-cysteine than wild type^7^. Here, overexpression of *cyuA* and *cyuP* alone did not produce H_2_S from GSH and was not significantly different from control (0 ± 0 μM vs. 17 ± 25 μM, *p*>0.05, **Fig. 3b**). In contrast, overexpression of *ggt* and *pepB* enabled degradation of GSH to H_2_S (289 ± 96 μM). Co-expression of *ggt*, *pepB*, *cyuA*, and *cyuP* further increased H_2_S production (475 ± 132 μM), although this increase was not significant relative to the ggt-pepB strain. Finally, addition of the peptidase *pepT* did not further elevate H_2_S levels significantly (486 ± 146 μM).

**Fig. 3.**
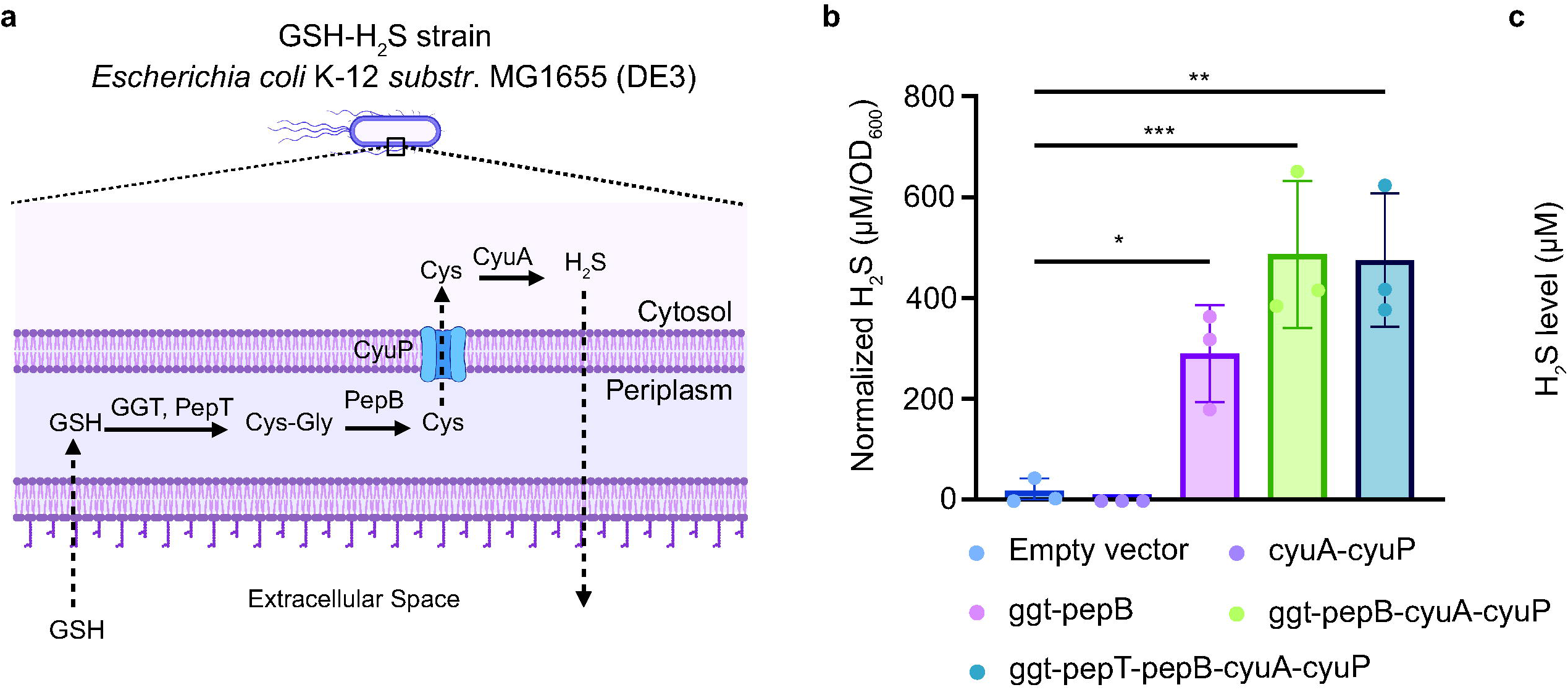
Engineered bacteria for producing H_2_S from glutathione in the lower gut. **a)** Schematic of GSH-H_2_S strain. GSH is converted to L-cysteinylglycine by *GGT* and *PepT*, and *PepB* cleaves L-cysteinylglycine to glycine and L-cysteine in the periplasm. L-cysteine is transported to the cytoplasm by *CyuP* and the L-cysteine desulfidase *CyuA* convert L-cysteine to H_2_S, which passively diffuses out of the cell. **b)** Cells were mixed in M9 media aerobically with 1 mM GSH for 3 hours. H_2_S was measured with the methylene blue assay and normalized to cell optical density (OD_600_). **c)** Engineered cells or empty vector cells were mixed in M9 media anaerobically in a 1:1 cell density ratio with five unique human fecal cultures. 1 mM GSH was supplemented and H_2_S was measured after 3 hours with the methylene blue assay. Error bars represent SD, and bars represent the mean value. *p<0.05, ∗∗p < 0.01,***p<0.001 ∗∗∗∗p < 0.0001. Significance was determined with an unpaired t-test with Welch’s correction or one-way ANOVA with post hoc Tukey analysis, with statistical significance set at p < 0.05. Figures made with Biorender.com

Having demonstrated H_2_S production from GSH in monoculture, we hypothesized the engineered microbe could produce H_2_S in the presence of the five human consortia, due to the low competition for GSH (**Fig. 1**). To test this, we used the *ggt*-*pepT*-*pepB*-*cyuA-cyuP* strain in a 1:1 ratio with each of the five human-derived consortia under anaerobic conditions. Excitingly, this strain produced significant levels of sulfide compared to the empty vector strain in co-culture minimal media supplemented with 1 mM GSH after three hours of anaerobic incubation (608 ± 90 μM vs. 28 ± 38 μM, *p*<0.0001) (**Fig. 3c**). The engineered phenotype is detectable across a panel of five heterogeneous human donors rather than in a single community.

When testing the strain in LB media, we noticed delayed onset of H_2_S production compared to minimal medium. We considered that peptides present in the peptone of LB might compete with GSH, either for transport into the engineered bacterium or for peptidase active sites, potentially limiting H_2_S production. To test this hypothesis, peptone was supplemented to minimal media and demonstrated partial suppression of H_2_S output compared to minimal media only (**Supp. Fig 4a**, 116 ± 3 μM vs. 475 ± 116 μM, p<0.01). Additionally, we explored heterologous expression of *fupA*, a porin from *Francisella tularensis* known to mediate GSH uptake, in *E. coli*, but this did not improve H_2_S production (**Supp. Fig. 4a**)^39^. The strain was tested in small intestinal extract, and minimal H_2_S production was observed relative to peptide-free media (**Supp. Fig. 4b**). The *ggt*-*pepT*-*pepB*-*cyuA-cyuP* strain suffers from peptide-mediated H_2_S production suppression, and peptides are rich in the small intestine. Based on these data, the GSH strain seems an ideal candidate for H_2_S delivery in the distal colon, as GSH is relatively stable in the colonic microbiota (**Fig. 1**), and the low concentration of peptides in the large intestine would not interfere with GSH uptake (**Supp. Fig. 4**)^29^.

### 3.4 Engineered bacteria assimilate H_2_S into glutathione anaerobically

Based on the low conversion of GSH to H_2_S by the human gut microbiota (**Fig. 1**), we hypothesized that GSH would also be a good sink molecule for sequestering intestinal sulfide. To test this, we designed two pathways to convert H_2_S into GSH (**Fig. 4a,b**). We overexpressed the native cysteine biosynthesis pathway, in which serine is first acetylated by a feedback-insensitive mutant of CysE (encoded by *cysE\*\**)^40^ to form O-acetylserine using acetyl-CoA, followed by nucleophilic sulfide incorporation catalyzed by CysK to produce L-cysteine from H_2_S. For conversion of L-cysteine to GSH, we co-expressed the native glutamate-L-cysteine ligase (*gshA*) and glutathione synthetase (*gshB)*, or the heterologous bifunctional glutathione synthetase (*gshF*) from *Streptococcus thermophilus* (**Fig. 4a,b**). The engineered strains were tested for sulfide depletion compared to empty-vector controls anaerobically in Hungate tubes, supplemented with serine, glycine, and glutamate. The initial *gshAB*- and *gshF*-expressing strains did not improve H_2_S degradation compared to control (**Fig. 4c**).

**Fig. 4.**
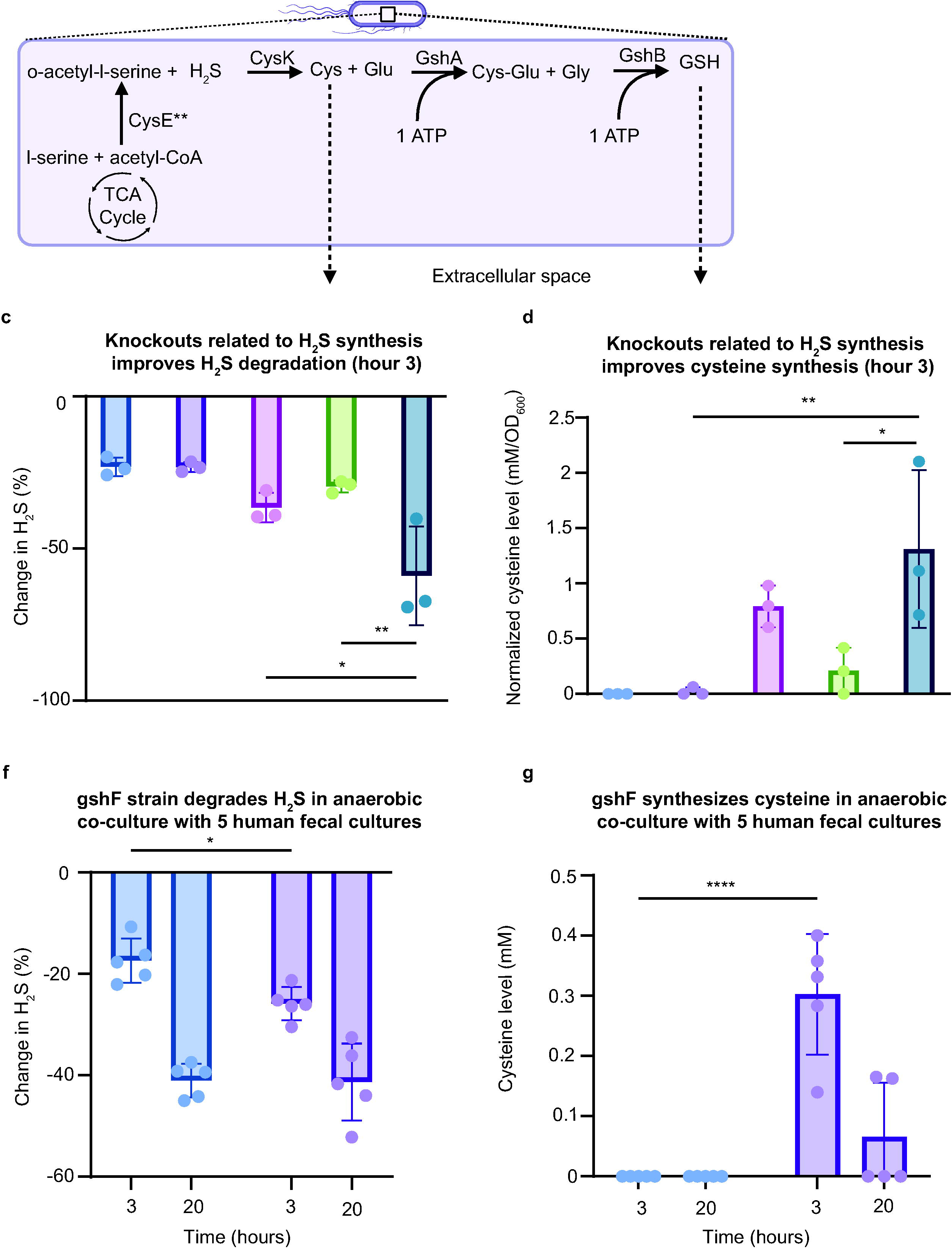
Engineered bacteria convert H_2_S to L-cysteine and glutathione. a,b) Metabolic pathways of gshAB and gshF strains. H_2_S is converted to L-cysteine and ligated to glycine and glutamate to make GSH. gshAB strain co-expresses the native *gshA* and *gshB* from *E. coli*, whereas gshF strain expresses the *S. thermophilus* bifunctional glutathione synthetase, *gshF*. **c)** H_2_S degradation under anaerobic conditions. **d)** L-cysteine synthesis under anaerobic conditions. **e)** GSH synthesis under anaerobic conditions. gshF (Quad KO) or empty vector control (Quad KO) strains were mixed with five human fecal donors in a 100:1 cell density ratio under anaerobic conditions, glycine, glutamate, and serine were supplemented. **f)** H_2_S levels**, g)** L-cysteine levels, and **h)** GSH levels in the co-culture with five human cultures. n = 3 independent experiments for monoculture experiments, n = 5 independent experiments for human fecal culture experiments. Error bars represent SD, and bars represent the mean value. *p<0.05, ∗∗p < 0.01, ∗∗∗∗p < 0.0001. One-way and two-way ANOVA with post hoc Tukey analysis, with statistical significance set at p < 0.05.”Figures made with Biorender.com

We hypothesized that the limited sequestration of sulfide might be due to the high activity of L-cysteine desulfidases, reconverting any L-cysteine formed back to sulfide in a futile cycle. To improve pathway flux towards GSH, we knocked out several L-cysteine desulfidases and related transcription factors alone or in combination, that reduced baseline sulfide production from L-cysteine. Specifically, we deleted: the *cyuAP* operon which contains a cysteine desulfidase and cysteine transporter; *cyuR*, the regulator of the *cyuAP* operon; and *iscS* which also has desulfidase activity^41,42^. Knockout strains were tested for changes in growth rates and showed trending decreases in H_2_S production from L-cysteine (**Supp. Fig. 5,6**). The quadruple knockout (MG1655-Δ*cyuR-cyuP-cyuA-iscS*, named “Quad KO”) was used as the genetic background for subsequent engineering and compared to WT background for improved H_2_S conversion to L-cysteine and GSH.

The Quad KO strain expressing *gshF* significantly reduced H_2_S levels relative to both the corresponding WT control, and the Quad KO strain expressing *gshAB* strains after 3 hours of anaerobic culture with sulfide (**Fig. 4c**). This strain also increased extracellular L-cysteine levels compared with its WT background at all time points (**Fig. 4d, Supp. Fig. 7**). Both gshAB (Quad KO) and gshF (Quad KO) strains elevated extracellular GSH levels compared to their respective WT background strains at all time points (**Fig. 4e, Supp. Fig. 7**). Collectively, deletion of these four H_2_S-related genes enhanced conversion of H_2_S into L-cysteine and GSH. Based on its superior H_2_S degradation capacity, particularly under anaerobic conditions, the gshF-expressing Quad KO strain was selected as the lead candidate.

To test its ability to convert H_2_S in a more complex environment, we next mixed the gshF Quad KO strain with the human consortia under anaerobic conditions. This strain significantly lowered H_2_S compared to an empty vector control after three hours (-26 ± 3% vs. -17 ± 4%, p<0.05), but not at 20 hours (**Fig. 4f**). This is likely due to the exported L-cysteine that accumulates in the extracellular media (**Fig. 4g**), which can be rapidly converted by the microbiota back to H_2_S (**Fig. 1a,b)**. GSH levels trended higher with the GSH Quad KO strain, though this observation was not statistically significant (**Fig. 4h**, *p*=0.0638). We therefore turned our attention to alternative pathways and sulfur sinks.

### 3.5 An Sqr-expressing microbe improves H_2_S degradation kinetics in human fecal consortia

The gshF-based H_2_S consumption strain exhibited several limitations that constrained its performance in complex microbial communities. The glutathione pathway requires three amino acids and intracellular ATP to synthesize GSH. Additionally, flux imbalance from poor catalytic efficiency of GshF results in accumulation and exports the pathway intermediate, L-cysteine, which can be rapidly converted back to H_2_S by the microbiota. This motivated our search for an alternative pathway with reduced metabolic demands, faster kinetics, and a stable sulfur byproduct. Sulfide:quinone reductase (Sqr) represents an attractive alternative for microbial H_2_S consumption, and human fecal consortium data suggest its potential activity in the gut microbiome (**Fig. 1**). Sqr enzymes generally exhibit k_cat_/K_m_ values several orders of magnitude higher than glutathione synthetases (**Table S5**), suggesting superior catalytic efficiency^43^. We previously demonstrated that Sqr from *Wolinella succinogenes* functions in *E. coli* under both aerobic and anaerobic conditions, converting H_2_S to persulfide products^32^. It has also been shown that Sqr oxidizes sulfide to octasulfur clusters in *E. coli*^44^. We utilized the *W. succinogenes sqr*-expressing *E. coli* strain for improved sulfide degradation capacity (named “Sqr” strain, **Fig. 5a**).

**Fig. 5.**
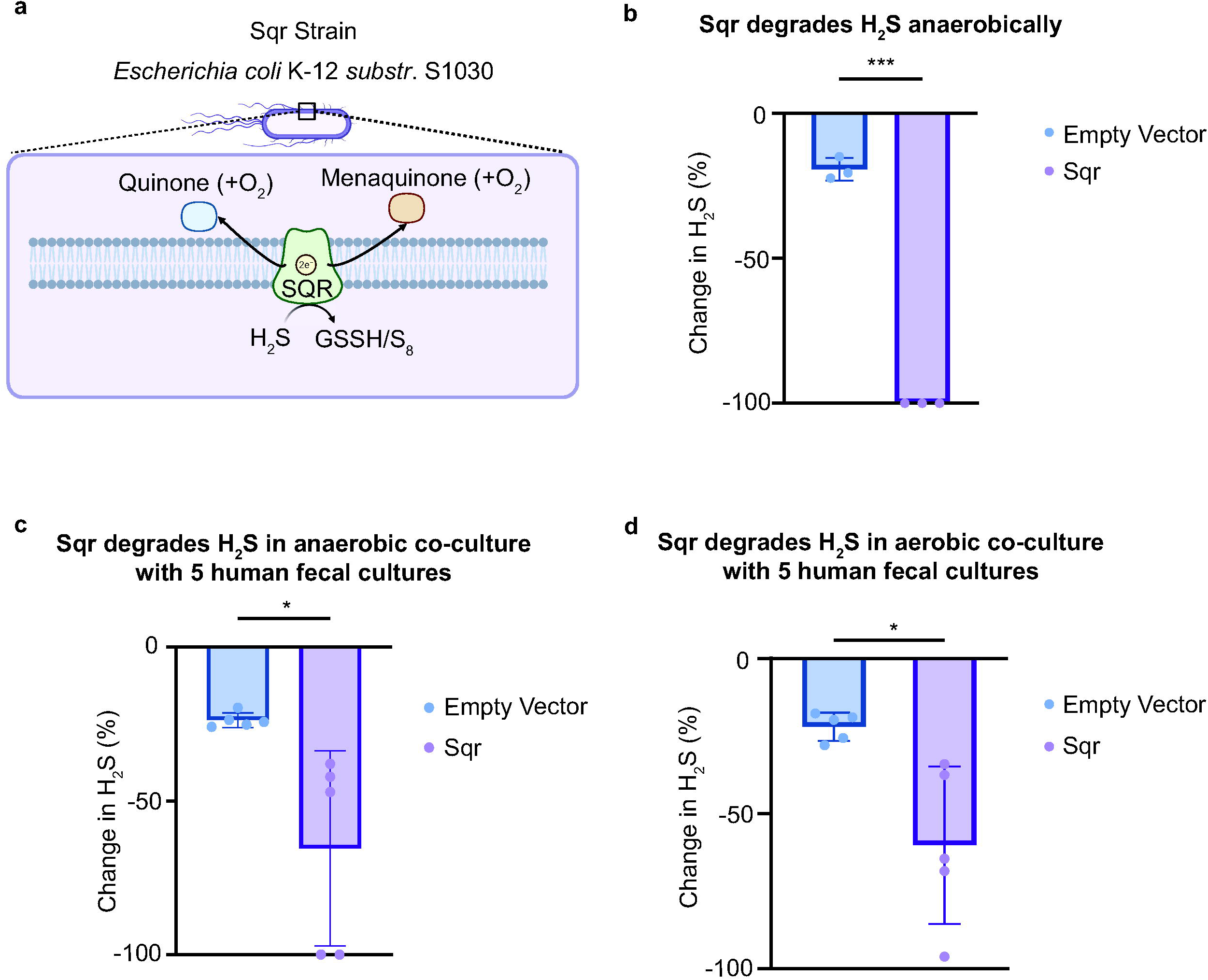
Sqr-expressing bacteria degrade H_2_S superior to glutathione synthesis strain. **a)** Schematic of Sqr strain. Sqr oxidizes sulfide, passing two electrons to quinone (aerobic conditions) or menaquinone (anaerobic conditions). The sulfur atom is integrated into a persulfide (GSSH) or into octasulfur clusters (S_8_). **b)** Sqr and EV strains were mixed with ∼1.5 mM Na_2_S in anaerobic media supplemented with fumarate and nitrate in Hungate tubes, H_2_S was sampled after three hours with the methylene blue assay. **c)** Sqr or EV strains were mixed with five human fecal donors in a 10:1 cell density ratio under anaerobic conditions supplemented with nitrate and fumarate, H_2_S was sampled after three hours. **d)** Sqr or EV strains were mixed with five human fecal donors in a 1:1 cell density ratio under aerobic conditions, H_2_S was sampled after three hours. n = 3 independent experiments. Error bars represent SD, and bars represent the mean value. *p<0.05, ***p<0.001. Significance was determined with an unpaired t-test with Welch’s correction, with statistical significance set at p < 0.05.Figures made with Biorender.com

In anaerobic monoculture supplemented with fumarate and nitrate as terminal electron acceptors, the Sqr strain depleted 100% of supplied H_2_S within 3 hours, significantly outperforming the empty vector control (100 ± 0% vs. 19 ± 4%, p<0.001) (**Fig. 5b**). To assess performance under more physiologically relevant conditions, we co-cultured the Sqr strain with human fecal microbiota at various ratios under anaerobic conditions. At a 10:1 ratio, the Sqr strain achieved significantly greater H_2_S degradation than the empty vector control (-65 ± 32% vs. -24 ± 2%, p<0.05) (**Fig. 5c**), though performance at a 1:1 ratio showed a non-significant trend toward reduced H_2_S levels (**Supp. Fig. 8**). Given that healthy and inflamed intestinal tissues maintain microaerobic conditions at the mucosal interface^45^, we also evaluated the capacity of the Sqr strain to utilize oxygen as a terminal electron acceptor in co-culture with human fecal microbiota at a 1:1 cell density ratio. The Sqr strain oxidized significantly more H_2_S than the empty vector control after 3 hours (-60 ± 25% vs. -22 ± 5%, p<0.05) (**Fig. 5d**). These data demonstrate that the *sqr-*expressing strain has metabolic versatility across oxygen tensions relevant in different intestinal regions and can degrade sulfide in the presence of human fecal gut microbiota.

To test the Sqr strain’s ability to lower H_2_S along the GI *in vivo*, it was necessary to artificially elevate intestinal sulfide levels first, as healthy mice have relatively low levels of intestinal H_2_S (**Fig. 2b-e**). The L-cysteine-H_2_S producing strain was used to artificially elevate sulfide levels and test the capacity of the Sqr strain to degrade sulfide *in vivo*. We gavaged the Sqr or empty vector strain with fumarate, and the L-cysteine-H_2_S strain and L-cysteine (**Supp. Fig. 9**). In both experiments, the H_2_S-producing strain increased luminal H_2_S levels. For the Sqr and empty vector groups, luminal H_2_S levels were compared. Sqr did not lower intestinal sulfide levels, but trended towards decreases in the stomach. Similarly, we tried using GYY4137 to elevate sulfide for testing Sqr, though its ability to reduce sulfide levels were insignificant (**Supp. Fig. 10**). The Sqr strain was tested *ex vivo* in small intestinal extract by supplementing exogenous sulfide, and significantly reduced sulfide levels relative to an empty vector control (**Supp. Fig. 11**). These data show Sqr functions in complex *ex vivo* environments, though establishing an *in vivo* elevated H_2_S model and measuring Sqr activity is challenging and could be the subject of future work.

## 4. Discussion

H_2_S is short-lived and chemically reactive, making controlled delivery difficult. Small molecule donors including GYY4137, allyl polysulfides, and sulfide salts have shown promise, yet their H_2_S-release kinetics can be pH-dependent and difficult to predict under physiological conditions^15^. By engineering *E. coli* sulfur metabolism, we enabled programmable control over H_2_S levels in both *ex vivo* human fecal microbiota cultures and animal models. Our findings establish that engineered bacteria can function as living delivery systems for gaseous H_2_S to the mammalian gut, with potential use in both fundamental biological discovery and medical applications.

We investigated native gut microbiota sulfur metabolism to inform strain design and guide selection of sulfur substrates and sinks. GSH was poorly converted to H_2_S relative to sulfite, thiosulfate, and L-cysteine. In aerobic metabolism, GSH donates electrons to reactive oxygen species (ROS), limiting oxidative stress in the cell. In anaerobic metabolism, ROS levels are much lower, and some anaerobes use alternative low-molecular-weight thiols, such as mycothiol or bacillithiol, to maintain redox balance and detoxify reactive compounds. This makes GSH a prime target as a sulfur sink or source in the anaerobic gut.

The microbiota coupled sulfide oxidation to nitrate reduction, though not fumarate, suggesting the potential role of *Sqr* to lower extracellular H_2_S levels, which is supported by metagenomics data showing the presence of the gene in the human gut microbiome^27^. We integrated these metabolic insights with aspects of human intestinal physiology, including regional variations in pH, nutrient availability, microbial density, and oxygen tension for localized H_2_S modulation for specific gastrointestinal regions^29^.

The contribution of the small intestinal microbiota to both intestinal and systemic sulfide levels is increasingly recognized^47^. Historically, its impact was underappreciated compared with the colonic microbiota, largely because sampling the small intestine is more invasive than collecting fecal samples. The L-cysteine-H_2_S producer was designed to target the upper gastrointestinal tract. The strain produces H₂S rapidly, allowing it to act within the short residence time of the stomach and small intestine (2-8 hours total in humans). Its activity is not known to be affected by competing dietary substrates, such as peptides, and the relatively low microbial abundance in the small intestine reduces L-cysteine competition (duodenum: 10^3^ CFU/mL; jejunum: 10^4^-10^5^ CFU/mL; ileum: 10^7^-10^8^ CFU/mL)^29^. To limit sulfide production to the upper gut, L-cysteine was selected as the sulfur source due to its rapid absorption in the small intestine. The fate of the amino acid is 1) conversion to H_2_S by the engineered strain, or 2) absorption in the small intestine via the SLC7A11 L-cysteine/glutamate antiporter^48,49^. Previous research showed >90% of dietary L-cysteine is absorbed in the small intestine in both mice and humans^48,50^. In agreement with previous work, we measured nearly zero free L-cysteine in the cecum and colon of mice following gavage with high concentrations of the amino acid and the control bacteria (**Supp. Fig. 3**).

In our mouse studies, the L-cysteine-based strain elevated H_2_S along the gastrointestinal tract. We also gavaged GYY4137, the gold-standard H_2_S-releasing molecule, which only elevated levels in the stomach. GYY4137 releases H_2_S in a pH-dependent manner, with accelerated release at lower pH, limiting precise control along the pH-variable gut^37^. Although sodium bicarbonate was added to oral gavages, this only temporarily neutralizes stomach acid, which can re-acidify within 30 minutes to release sulfide from GYY4137^51^. GYY4137 did not elevate small intestinal levels compared to control, likely due to mildly acidic pH (pH 5-6), compared to lower acidity of the stomach.

In contrast, the GSH-based H_2_S producer is designed for the lower gastrointestinal tract, exploiting its unique metabolic capacity to utilize GSH as a substrate for H_2_S production anaerobically. We showed this pathway is largely inactive in native colonic microbiota. However due to limitations of gavaging GSH, specifically rapid degradation and absorption of GSH in the small intestine via peptidases and amino acid transporters^52^, this strain was not evaluated *in vivo*. In humans and complex pre-clinical models, delivery of probiotics and payloads to the colon is feasible by use of enteric-coated capsules, which could be leveraged to test the strain’s ability to elevate sulfide levels in the lower gut^53^. However, size constraints for enteric coated tablets have rendered these techniques unreliable in rats and nearly impossible in mice^54^. These strategies highlight the merging of metabolic engineering and human gut physiology to create more location-specific probiotics.

The H_2_S degrading, GSH-producing strain exhibited poor sulfide degradation under anaerobic conditions in Hungate tubes. Through deletion of genes involved in L-cysteine catabolism, we significantly enhanced H_2_S degradation capacity and increased L-cysteine and GSH synthesis. Further optimization using a heterologous glutathione synthetase improved these metrics. However, the high metabolic demand (i.e., amino acids, ATP, acetyl-CoA) and slow reaction kinetics of this strain fragilizes it to the metabolic heterogeneity of the gut. This was shown by its poor H_2_S degradation capacity and high extracellular L-cysteine accumulation in the human fecal microbiota cultures. To improve sulfide degradation, we tested the Sqr strain which offers versatility for both upper and lower gut applications. It can use oxygen, fumarate, or nitrate as electron acceptors, does not require amino acids or ATP for sulfide oxidation, and produces persulfides or octasulfur, sequestered intracellularly, as a byproduct^32,44^. The Sqr strain’s H_2_S degradation kinetics were superior to the GSH strain in both anaerobic monoculture and *ex vivo* human fecal microbiota. This is in line with prior work showing k_cat_/K_m_ values for *Sqr* are typically superior to those for glutathione synthetases (**Table S5**).

To address low basal GI H_2_S levels, we used the L-cysteine-degrading, H_2_S-producing microbe, allowing us to test whether the Sqr strain could reduce luminal H_2_S compared with the empty vector strain. While Sqr may have converted H_2_S to persulfides or octasulfur *in vivo*, luminal H_2_S levels did not change. Direct measurement of these compounds via LCMS was unreliable as there was a high degree of background noise in luminal content. Additionally, there were several challenges with the experimental design. Effective *in vivo* H_2_S modulation requires spatial and temporal co-localization of the H_2_S-producing bacterium with L-cysteine and of the Sqr-expressing strain with its electron acceptor fumarate. Further, the short nature of the experiments (1-hour), potentially limited the window of activity for the Sqr strain. Longer-term studies with stable intestinal colonization or daily gavaging would better model medical or chronic disease scenarios. Colonization of the intestinal tract with H_2_S-producing strains, particularly when coupled with dietary L-cysteine supplementation, could enable sustained elevation of H_2_S levels over long durations, as we previously showed in human gut-on-chip systems, for investigating its role in IBD, IBS, or SIBO^7^. Physiological read-outs such as thiosulfate and inflammatory markers could give insight into the impacts of sulfide on host metabolism, while measuring microbiota composition and gene expression could reveal its impact on the microbiome.

Reported colonic sulfide concentrations span a wide range (0.2-3.4mM), in part because total and free sulfide differ by up to two orders of magnitude: the majority of sulfide is bound as acid-labile sulfide, and only the free pool is bioavailable to the epithelium^46^. More recent analysis suggests the free colonic sulfide range is 10-190 μM in humans, and 2-120 μM in rodents. Most analyses, however, rely on *ex vivo* culturing of fecal samples and measurement of the H_2_S generated, an approach that fails to capture luminal H_2_S. In this study, we instead measure free sulfide directly in the gut by rapid derivatization with mBBr, capturing a snapshot in time.

In summary, we developed engineered strains to modulate sulfide levels, which could be used to decipher its contradictory role in disease. These strains also have strong translational potential in medical applications. Beyond H_2_S, our approach could be applied to other intestinal metabolites, expanding the potential of precision-engineered probiotics for modulating host-microbiome interactions. Looking forward, this work showcases a generalizable framework for combining metabolic engineering and gut physiology principles to build more robust probiotics.

## Contributions

J.A.H., A.N.K., R.K., and B.M.W. conceived the study and experimental design. J.A.H. is the primary author and conducted the experiments and collected and analyzed the data. B.B. assisted in collecting *in vivo* data. A.W.L., T.K., and W.G. assisted in molecular cloning, data collection, and instrument set up. M.T.F. assisted in molecular cloning and protocol development. P.S. and M.M. contributed to the design and analysis of fecal microbiome studies. All authors reviewed and approved the final document.

## Competing Interests

J.A.H, R.A.K., and B.M.W are listed as inventors of pending patents related to work in this article. J.A.H, R.A.K, and B.M.W are co-founders/employees at, and have equity in Concordance Therapeutics Inc. P.S. and M.M are co-founders/employees at, and have equity in Holobiome Inc. The remaining authors declare no competing interests.

## Acknowledgements.

We thank Northeastern DLAM for experimental assistance. This material is based upon work supported by the National Science Foundation Graduate Research Fellowship under grant number 1938052 to J.A.H., by the National Institute of Biomedical Imaging and Bioengineering of the National Institutes of Health under award number R21EB033892 to B.M.W, and we gratefully acknowledge support from Propel a Cure for Crohn’s Disease and the Mendez Family Foundation. Any opinions, findings, conclusions, or recommendations expressed in this material are those of the authors and do not necessarily reflect the views of the funders.

