## Supplementary Information for "Metabolic engineering of *Escherichia coli* to modulate hydrogen sulfide levels in the mammalian gut"

### Supplemental Information

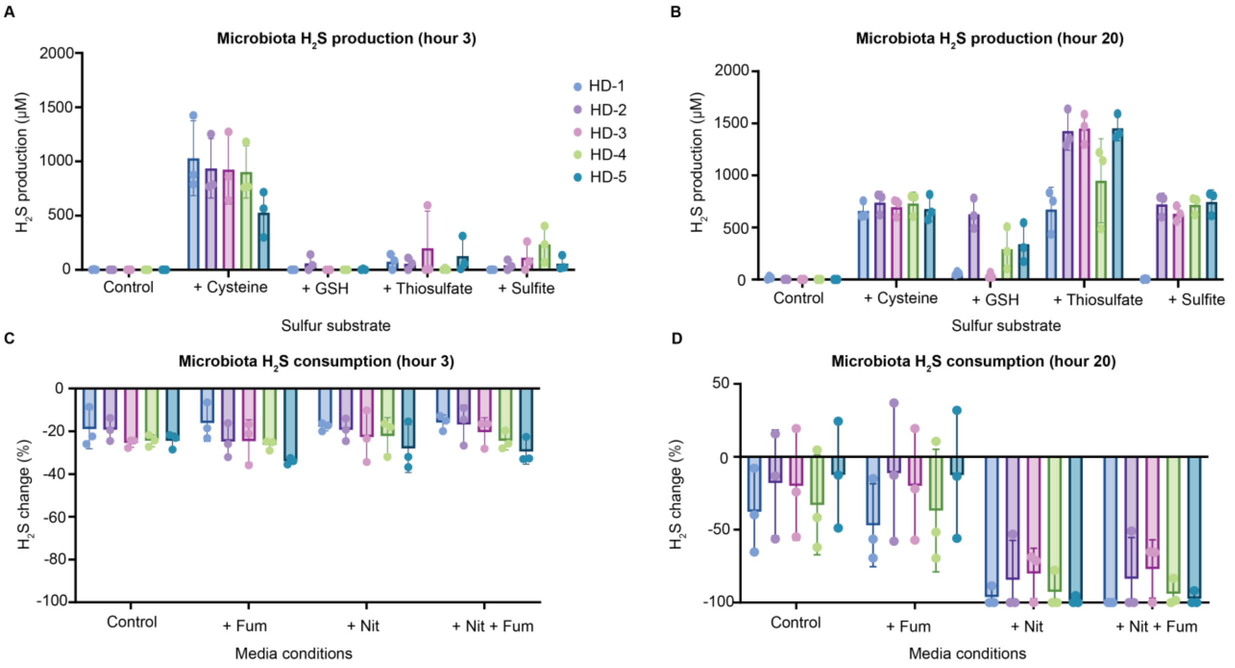


**Supp. Fig. 1 Human fecal microbiota produces and consumes $H_2S$. Related to Figure 1.** Five human fecal cultures were grown overnight anaerobically and resuspended in M9 minimal media and 1 mM of cysteine, glutathione, thiosulfate, or sulfite. Note, thiosulfate contains two sulfur atoms which can be reduced to sulfite, and sulfite to sulfide; values here are not normalized to sulfur added. $H_2S$ levels were measured via the methylene blue assay after **a)** 3 hours and **b)** 20 hours. Fecal consortia were prepared in the same manner and mixed with ~1.5 mM $Na_2S$ with or without electron acceptors, fumarate (Fum) and nitrate (Nit). $H_2S$ levels were measured after **c)** 3 hours and **d)** 20 hours. n = 3 independent experiments. Error bars represent SD.

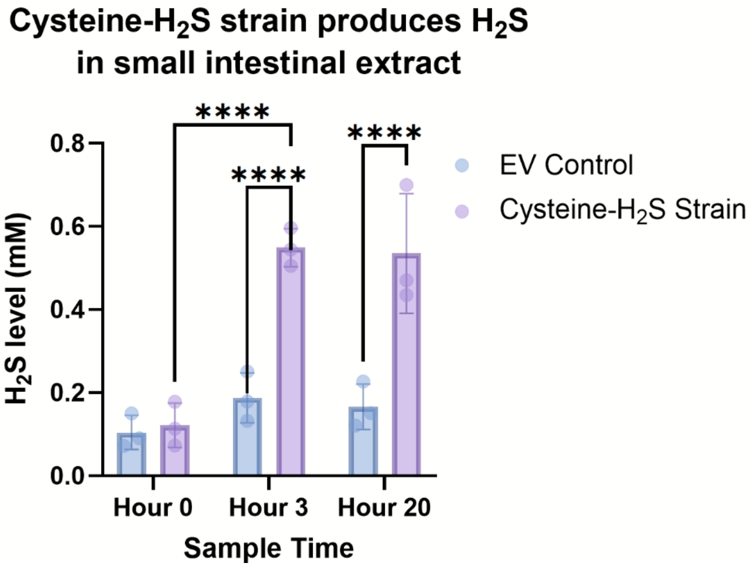


**Supp. Fig. 2 The cysteine-$H_2S$ strain is robust in small intestinal ex vivo model. Related to Figure 2.** Cells were prepared as described and mixed in PBS-small intestinal extract at $OD_{600}$ 1.0, and 1mM L-cysteine was added. Samples were taken with the mBBr assay to overcome challenges with the methylene blue assay in complex *ex vivo* samples. n = 3 independent experiments. Error bars represent SD, and bars represent the mean value. $**p < 0.01$, $****p < 0.0001$. Two-way ANOVA with post hoc Tukey analysis using a 95% confidence interval.

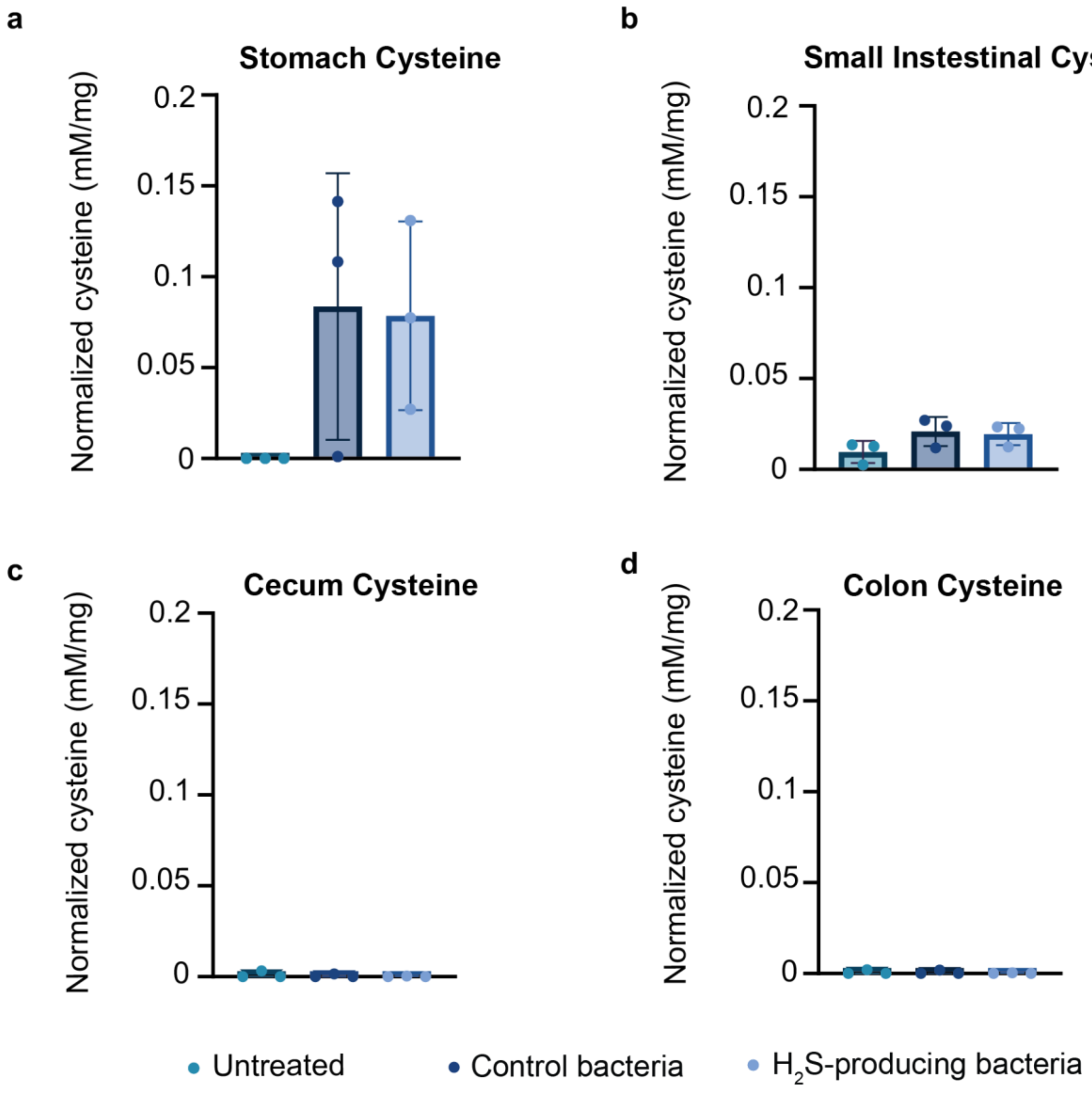


**Supp. Fig. 3. Intestinal cysteine levels from mouse experiments.** Untreated animals, animals treated the EV control strain and L-cysteine, or L-cysteine-$H_2S$ strain and L-cysteine were dissected and intestinal content assayed with mBBr. Cysteine levels were quantified as described. Error bars represent SD, and bars represent the mean value.

a

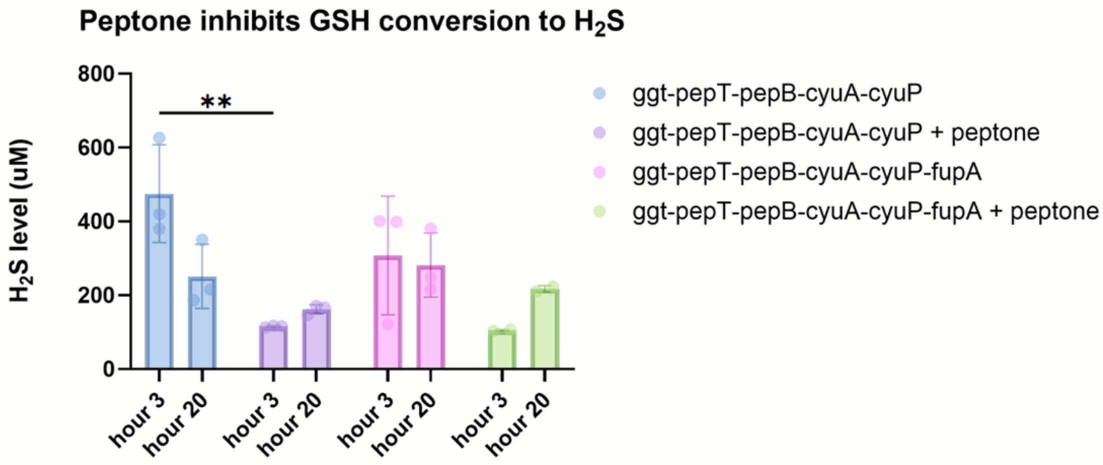


b

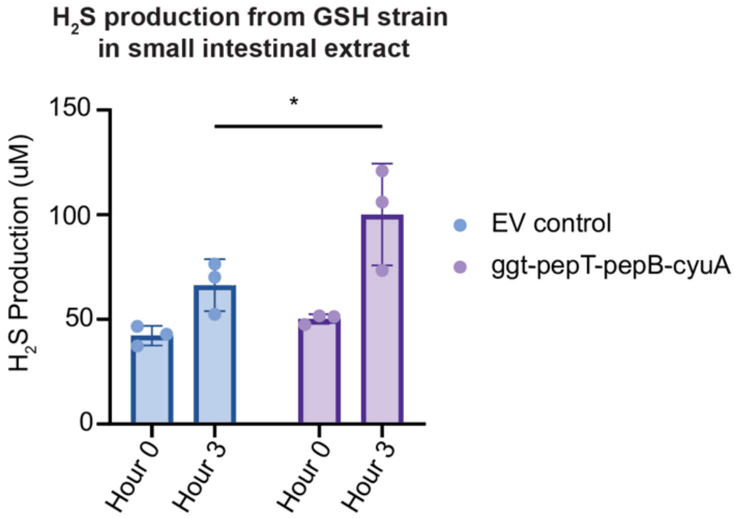


**Supp. Fig. 4 Peptone inhibits GSH-$H_2S$ strain. Related to Figure 3. a)** Cells were prepared as described and mixed in minimal M9 media with 1 mM GSH and 10g/L peptone. $H_2S$ was measured with the methylene blue assay. **b)** Cells were prepared as described and mixed in PBS-small intestinal extract at $OD_{600}$ 1.0, and 1mM GSH was added. Samples were taken with the mBBr assay to overcome challenges with the methylene blue assay in complex *ex vivo* samples. n = 3 independent experiments. Error bars represent SD, and bars represent the mean value. $**p < 0.01$, $****p < 0.0001$. Two-way ANOVA with post hoc Tukey analysis using a 95% confidence interval.

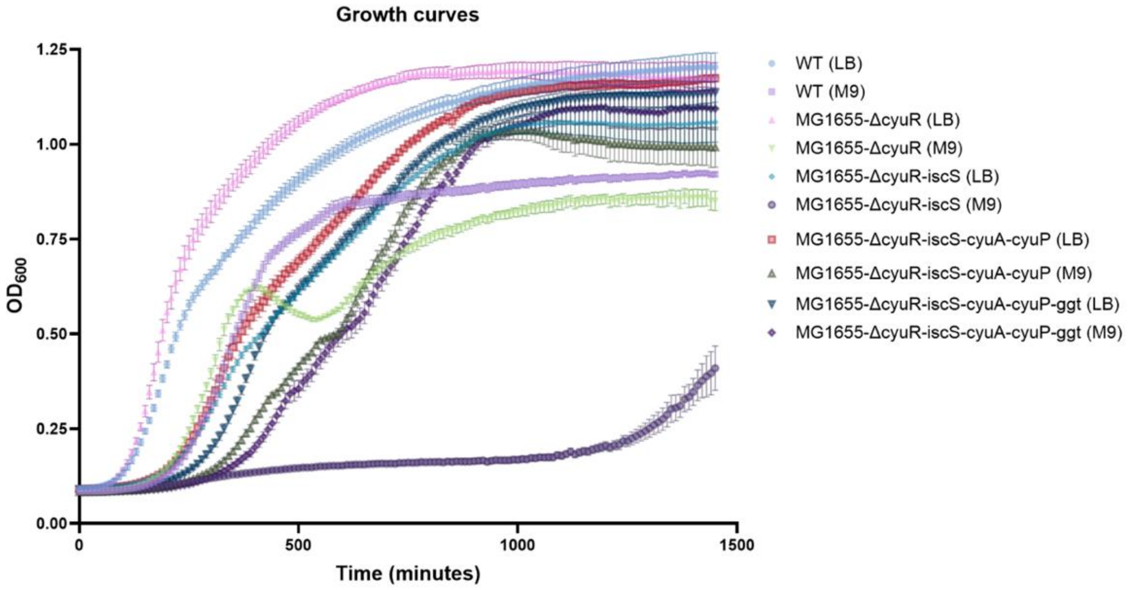


**Supp. Fig. 5 Growth curves associated with each genetic background. Related to Figure 3. a)** Cells were prepared as described and inoculated in 96-well plates at $OD_{600}$ 0.01 in either M9 minimal media or LB media in technical triplicates. $OD_{600}$ reads were taken every 10 minutes for 16 hours. Error bars represent SD, and solid lines represent the mean value.

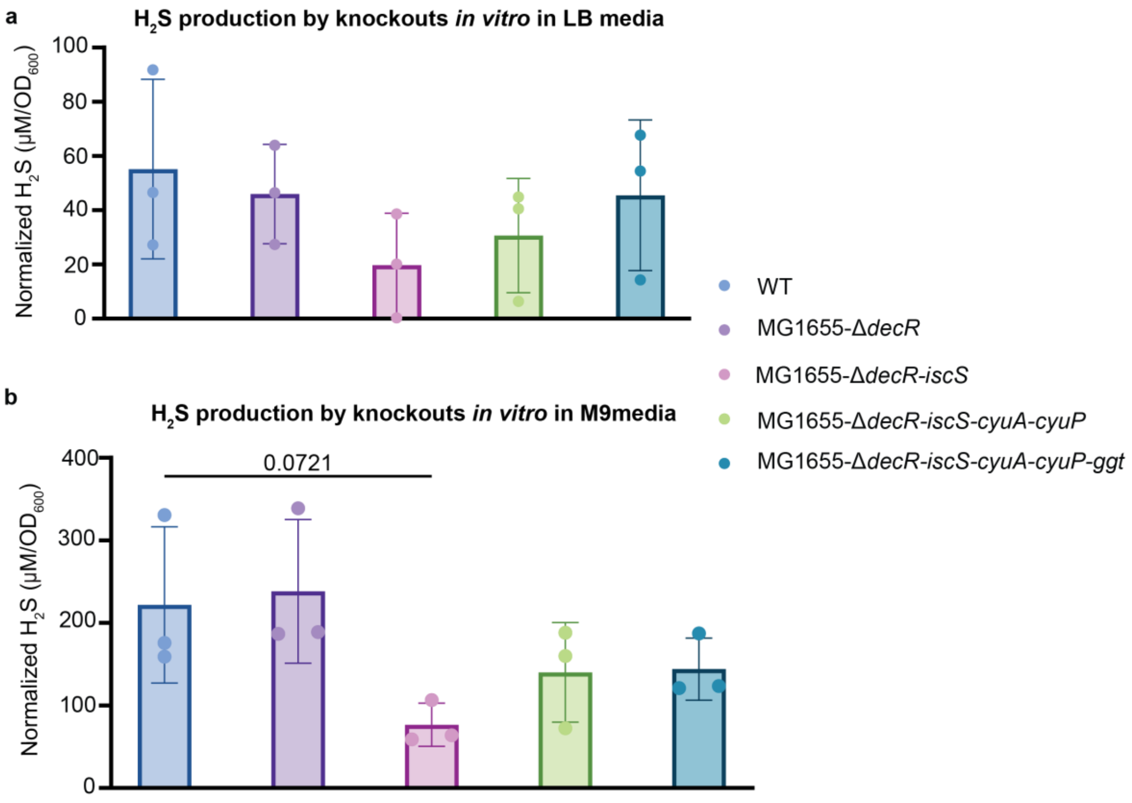


**Supp. Fig. 6 Knockouts related to $H_2S$ production trends towards lower $H_2S$ production from L-cysteine in M9 and LB media. Related to Figure 3.** Cells were prepared as described and mixed with 1mM L-cysteine in aerobic conditions in **a)** M9 media and **b)** LB media. $H_2S$ was measured with the methylene blue assay after three hours of culture. n = 3 independent experiments. Error bars represent SD, and bars represent the mean value. One-way ANOVA with post hoc Tukey analysis using a 95% confidence interval.

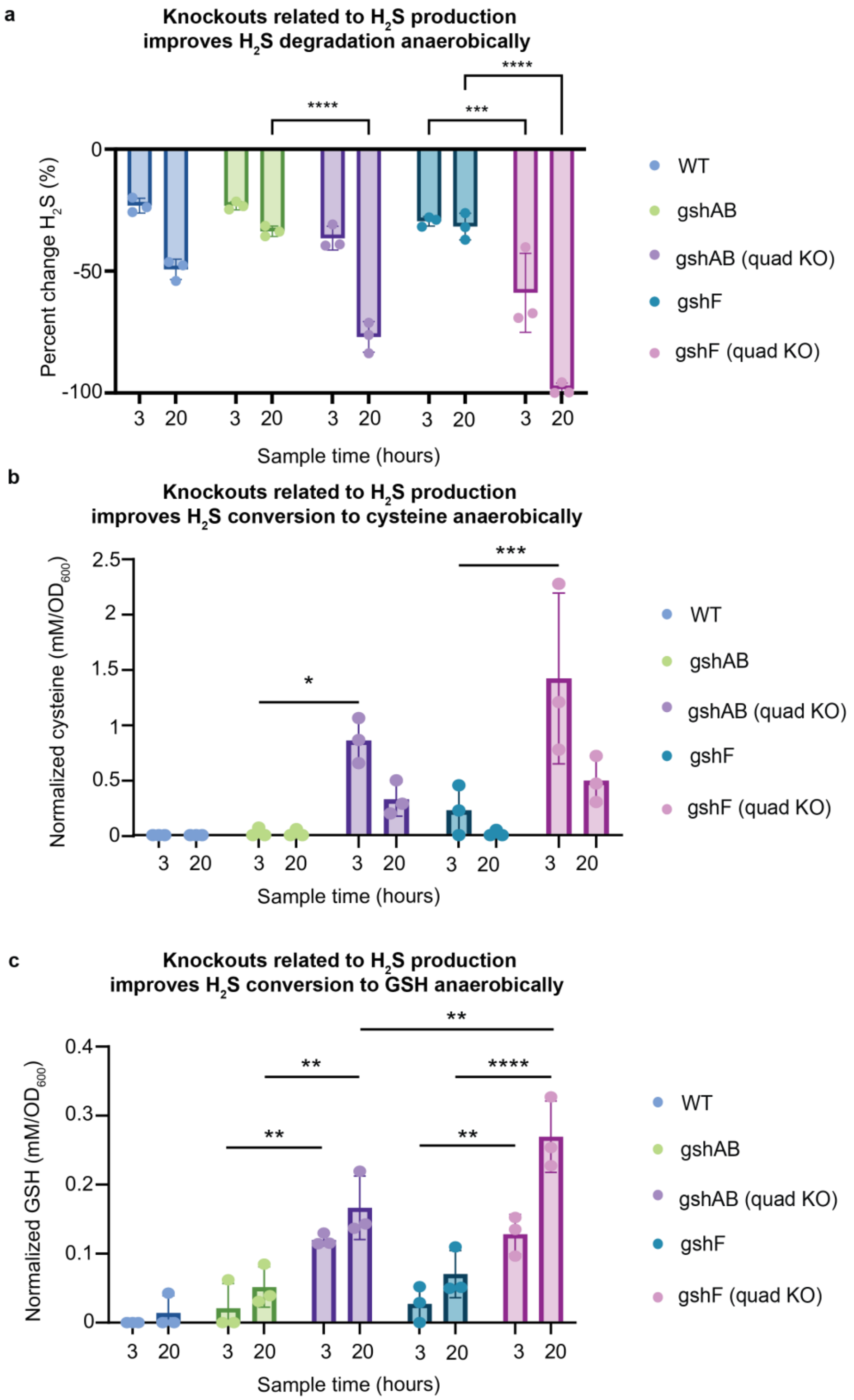


**Supp. Fig. 7 Engineered bacteria convert $H_2S$ to cysteine and glutathione. Related to Figure 3. a)** $H_2S$ degradation, **b)** cysteine synthesis, and **c)** glutathione synthesis by strains under anaerobic conditions with glycine, glutamate, and serine supplemented. n = 3 independent experiments. Error bars represent SD, and bars represent the mean value. *p<0.05, ⁎⁎p < 0.01,***p<0.001, ⁎⁎⁎⁎p < 0.0001. Two-way ANOVA with post hoc Tukey analysis using a 95% confidence interval.

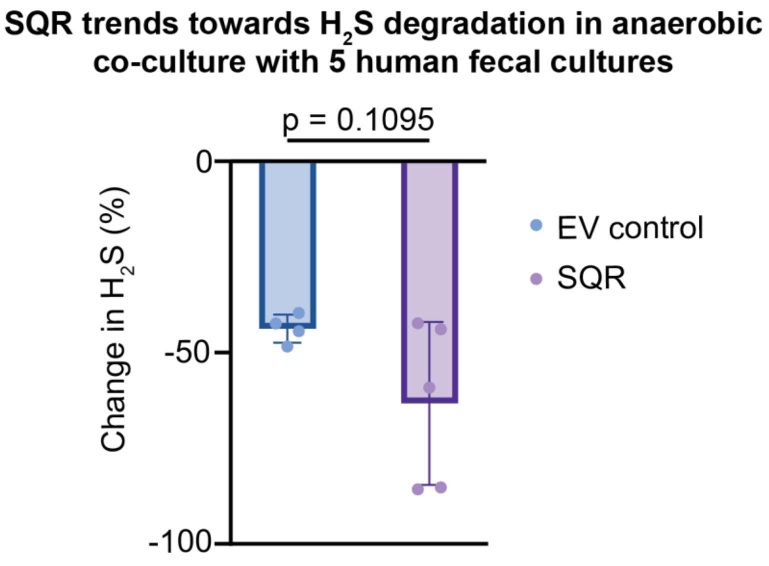


**Supp. Fig. 8. SQR strain trends towards decreased $H_2S$ levels in co-culture with 5 human fecal cultures under anaerobic conditions and 1:1 cell density. Related to Figure 4.** SQR or the EV control were mixed at a 1:1 cell density ratio ($OD_{600}$) in anaerobic M9 media with 1.5 mM Na2S, 20 mM sodium fumarate, and 20 mM sodium nitrate added. Sample taken after 3 hours of co-culture. n = 5 independent experiments. Error bars represent SD, and bars represent the mean value. Significance was determined with an unpaired t-test with Welch's correction using a 95% confidence interval.

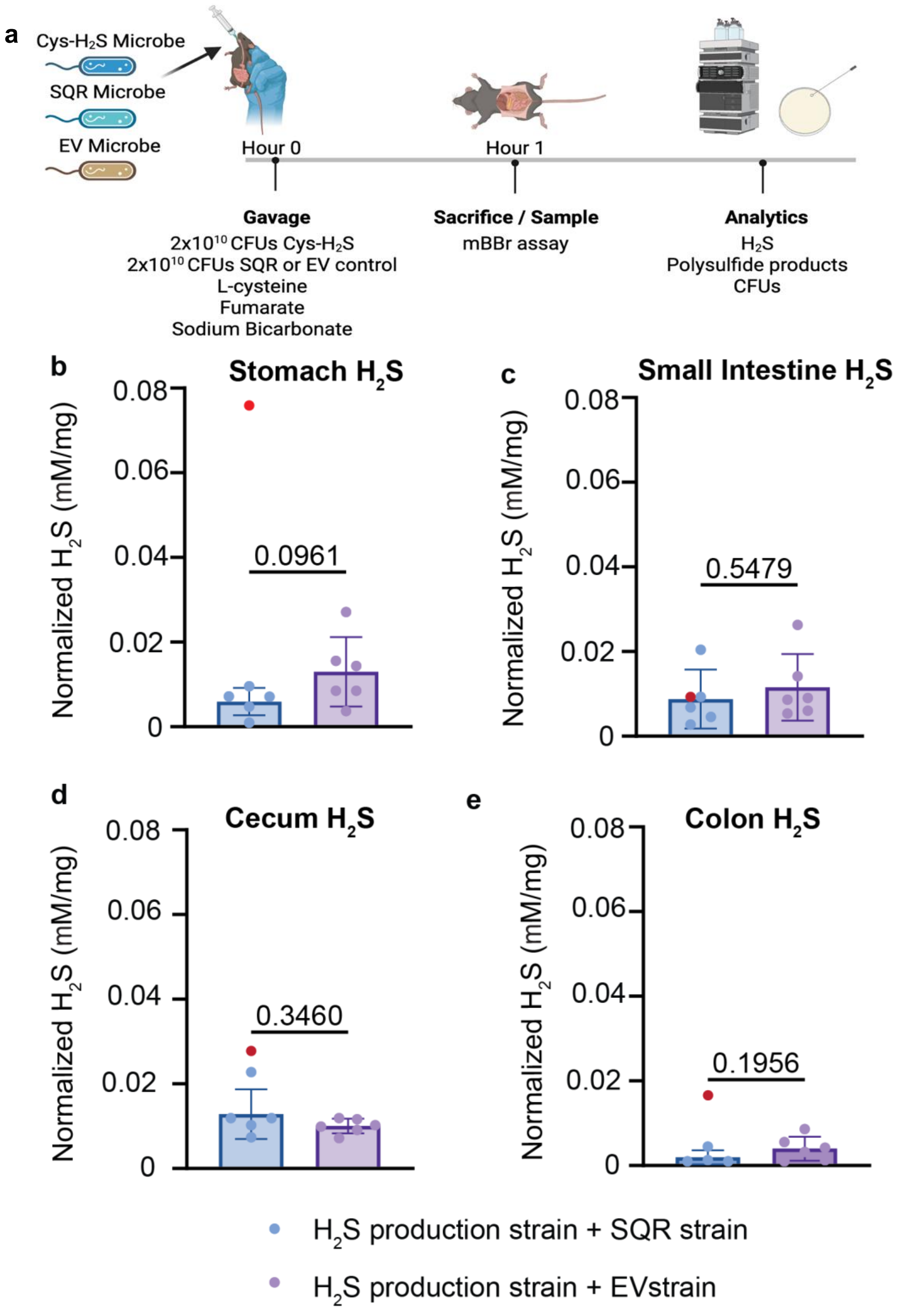


**Supp. Fig. 9 Intestinal $H_2S$ levels of mice co-gavaged with SQR or EV control, and $H_2S$-producing microbe.** Animals were sacrificed and luminal content mixed with the mBBr assay. $H_2S$ values normalized to sample weight in the **b)** stomach **c)** small intestine (S.I.), **d)** cecum, and **e)** colon. For EV Control n=6, 3 females, 3 males; SQR: n=6, 3 females, 3 males. Red data point represents the outlier (Z-score >21), and wasn't included in calculating the mean, standard deviation, or p value. Error bars represent SD, and bars represent the mean value. Significance determined by unpaired t test with Welch's correction.

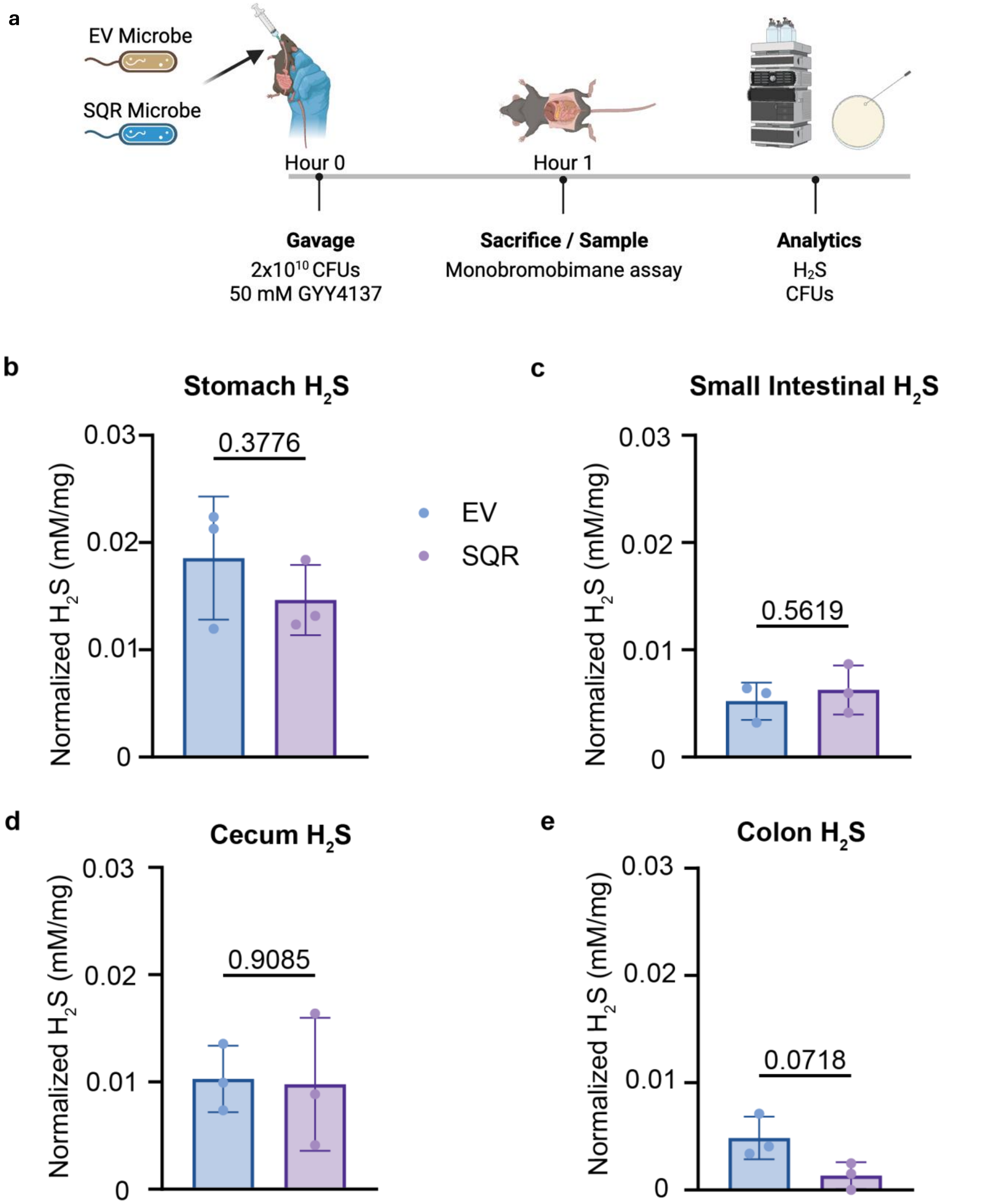


**Supp. Fig. 10 Intestinal $H_2S$ levels of mice gavaged with SQR or EV control and GYY4137. Related to Figure 6.** Animals were sacrificed and luminal content mixed with the mBBr assay. $H_2S$ values normalized to sample weight in the **b)** stomach **c)** small intestine (S.I.), **d)** cecum, and **e)** colon. For EV Control n=3, 2 females, 1 male; SQR: n=3, 1 female, 2 males. Error bars represent SD, and bars represent the mean value. Significance determined by unpaired t test with Welch's correction.

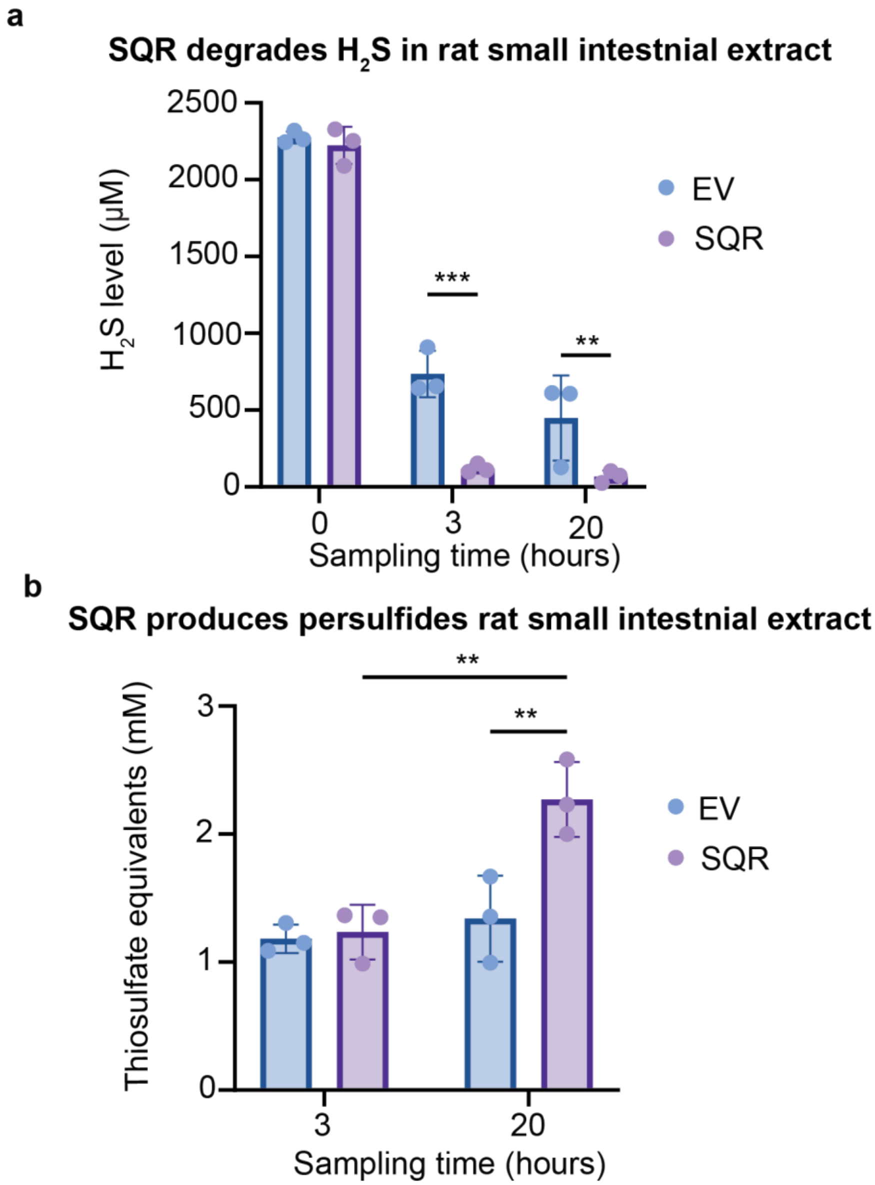


**Supp. Fig. 11 SQR strain degrades $H_2S$ in small intestinal extract from rats. Related to Figure 6.** Small intestinal extract media and cells were prepared as described. SQR or EV Control cells were mixed with 5 mL of extract media and supplemented with 2mM $Na_2S$ and levels were sampled at three timepoints. **a)** $H_2S$ levels were sampled with the mBBr assay and **b)** thiosulfate equivalents were determined using the cyanolysis assay, confirming persulfide synthesis by SQR. Error bars represent SD, and bars represent the mean value. $**p < 0.01$, $***p<0.001$. Two-way ANOVA with post hoc Tukey analysis using a 95% confidence interval.
